# The Amyloid Precursor Protein is a conserved Wnt receptor

**DOI:** 10.1101/2021.01.18.426557

**Authors:** Tengyuan Liu, Maya Nicolas, Tingting Zhang, Heather Rice, Alessia Soldano, Annelies Claeys, Iveta M. Petrova, Lee G. Fradkin, Bart De Strooper, Bassem A. Hassan

## Abstract

The Amyloid Precursor Protein (APP) and its homologues are transmembrane proteins required for various aspects of neuronal development and activity, whose molecular function is unknown. Specifically, it is unclear whether APP acts as a receptor, and if so what its ligand(s) may be. We show that APP binds the Wnt ligands Wnt3a and Wnt5a and that this binding regulates APP protein levels. Wnt3a binding promotes full length APP (flAPP) recycling and stability. In contrast, Wnt5a promotes APP targeting to lysosomal compartments and reduces flAPP levels. A conserved Cysteine Rich Domain (CRD) in the extracellular portion of APP is required for Wnt binding, and deletion of the CRD abrogates the effects of Wnts on flAPP levels and trafficking. Finally, loss of APP results in increased axonal and reduced dendritic growth of mouse embryonic primary cortical neurons. This phenotype can be cell-autonomously rescued by full length, but not CRD-deleted, APP.

## INTRODUCTION

The Amyloid Precursor Protein (APP) is the precursor that generates the Aβ peptide, whose accumulation is associated with Alzheimer’s disease (AD)[1]. As an ancient and highly conserved protein, APP and its homologs are found across animal species in both vertebrates and invertebrates[2]. As a result of the alternative splicing of the 18 exons coding for APP, there are 3 major isoforms expressed in different organs or tissues in mice and human[3]. APP695 is the major isoform expressed in the brain[4]. The expression of APP is detected at early stage during development[5,6]. In the developing mouse cortex APP mRNA is expressed continuously starting at embryonic day 9.5 coinciding with the initiation of neurogenesis and neuronal differentiation [7].

Structurally, APP is a type I transmembrane protein, which possesses a large extracellular amino acids sequence, an a-helix transmembrane sequence and a relatively short intracellular C-terminal sequence[8,9]. Based on the architecture of the ectodomain, APP has been proposed to be a putative receptor [10–14]. APP trafficking and processing have been intensively studied ever since the protein was first cloned. The turnover of transmembrane full length APP is rapid[15,16], and internalised APP can be degraded in lysosome or processed by α-, β- and y-secretase in different subcellular compartments to produce corresponding segments of APP. [17,18]. Recently, effort has been put into researching the function of the proteolytic products of APP under normal physiological condition[19], as this may provide new clues for AD research.

During *Drosophila* brain development the fly homolog of APP, called APPL, functions as key component of the neuronal Wnt-PCP signaling pathway and regulates the axonal outgrowth in fly mushroom body[20]. Both mammalian APP and fly APPL contain a Cysteine Rich Domain(CRD) in the ectodomain of APP which resembles the CRD of the binding domain of the Wnt Tyrosine-protein kinase receptor Ror2[9,21], suggesting the intriguing possibility that APP may itself be a receptor for Wnt family member.

Wnt signalling is an evolutionary conserved signal transduction pathway that regulates a large number of cellular processes. Three Wnt signaling pathway have been well described: the β-catenin based canonical pathway, the planar cell polarity (PCP/Wnt) signaling pathway and the calcium pathway. Wnt signaling regulates various features during development such as cell proliferation, migration and differentiation[22]. Recently, increasing evidence indicates that the Wnt signaling pathways are involved in the APP related Aβ production[23,24], but the precise mode of interaction between APP and the various Wnt pathways remains unclear.

The presence of CRD in APP, the reported involvement of Wnt signaling in APP processing and the importance of Wnt signaling during development suggested to us that APP may be a novel class of Wnt receptor regulating neuronal development. We used *Drosophila* and mouse embryonic primary cortical neurons as models to explore the APP-Wnt interactions during development. We provide evidence that the CRD of APP is a conserved binding domain for both canonical and PCP Wnt ligands. Furthermore, APP trafficking and expression is regulated by Wnts through the CRD, which in turn is required for APP to regulate axonal and dendritic growth and branching.

## RESULTS

### *Drosophila* APP Like interacts genetically with Wnt5

Drosophila APPL has been implicated in neural development[25,26] and is required for learning and memory [27]. *Drosophila* APPL shares high sequence homology with human APP and has been used as a model for understanding the physiological function of the APP family [28,29]. We previously reported that *appl* genetically interacts with components of the Wnt-PCP pathway [20] during mushroom body (MB) axon growth. The MBs are a bilateral neuronal structure in the fly brain required for learning and memory[30]. To understand the role of APPL in axonal PCP signaling, we first explored the specific nature of the genetic interaction between *appl* and the gene encoding the PCP protein Van Gough (Vang). In contrast to control MBs, 17% of male *appl* null mutant flies (*appl*^d^/Y, henceforth Appl−/−) displayed a loss of the MBb-lobe (Figure 1A, A’). The PCP receptor Vang is also required for β-lobe growth [31]; we observed that flies homozygous for the null allele *vang^stbm-6^* exhibited 50% β-lobe loss. Whereas *vang^stbm-6^* heterozygotes show no MB defects, the loss of one copy of *vang* in Appl−/− flies is comparable (43% β-lobe loss) to the complete loss of *vang* (Figure 1B). Therefore, in the absence of *appl, vang* is haploinsufficient. Next, we performed rescue experiments of Appl−/− mutant flies. Re-expression of APPL in the mutant MBs significantly rescued β-lobe loss. In contrast, the overexpression of Vang in Appl−/− null flies failed to do so. These loss and gain of function data suggest that Wnt-PCP signaling requires APPL to regulate axonal growth (Figure 1B).

**Figure 1:**
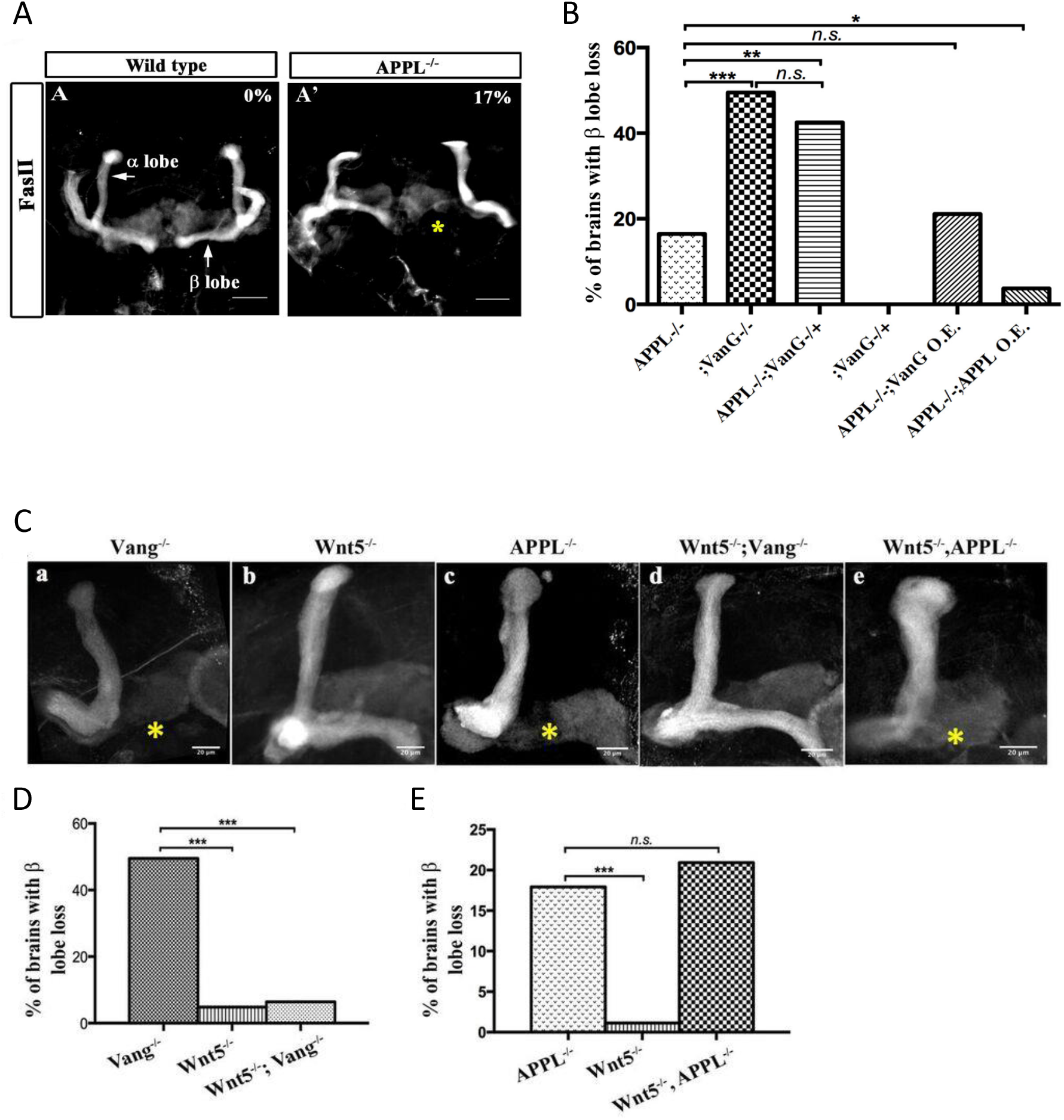
APPL mediates Wnt5a function in axonal growth. (A-A’) Structure of the MB neurons in adult wild type and APPL^−/−^ mutant flies. Immunofluorescence using anti-FascilinII (FasII) antibody that labels the axons of the MB. (A) In wild type brains, the axons of the MB project dorsally to form the *a* lobe and medially to form the β lobe. (A’) In APPL null mutant flies (APPL^d^/Y referred to as APPL^−/−^) there is axonal growth defect of the β lobe (as indicated by the asterisk) in 17% of the brains examined (n=97). Images are z-projections of confocal image stacks (scale bar, 50 μm). (B) APPL and VanG synergistically interact and APPL is necessary for VanG activity. The histogram shows the percentage of the β lobe defect in different genetic backgrounds. The loss of Vang induced a significantly higher penetrant phenotype up to 50%, (n=103) compared to APPL^−/−^; *p* value = 5.18^-7^ calculated with G-test. The loss of one copy of Vang in wild type background had no effect on axonal growth (n=30). However, the removal of one copy of Vang in APPL^−/−^ background significantly increased the phenotype to 43% (n=47) compared to APPL^−/−^; *p* value = 0.001026. The penetrance of the latter phenotype was not significantly different from the one observed in Vang^−/−^; *p* value = 0.4304. While the overexpression of APPL rescued the APPL^−/−^ phenotype (4%, n=54); *p* value = 0.01307, the overexpression of Vang failed to (21%, n=52); *p* value = 0.4901. * Indicates a *p* value<0.05. Data are shown as median ± whiskers. (Ca-e) Immunofluorescence analysis using anti-FasII antibody to show the structure of the MB axons in adult mutant flies of the following genotypes: (a) Vang^−/−^, (b) Wnt5^-^/Y referred to as Wnt5^−/−^, (c) APPL^−/−^, (d) Wnt5^−/−^,Vang^−/−^ and (e) Wnt5^−/−^,APPL^−/−^. Images are z-projections of confocal image stacks (scale bar, 20 μm). The asterisks correspond to the β lobe loss phenotype. (D) Wnt5 inhibits axonal growth, after branching, independently of Vang. The Histogram shows the percentage of the β lobe loss phenotype. Vang^−/−^ flies exhibit a highly penetrant phenotype of 50% (n=104), while Wnt5^−/−^ flies show a significantly less penetrant phenotype (5%, n=103); *p* value = 2.33^-14^ calculated with G-test. The loss of Wnt5 rescued Vang loss of function (6%, n=98); *p* value = 4.56^-12^. (E) Wnt5 inhibits axonal growth probably through APPL. Histogram showing the penetrance of the β lobe loss phenotype. In APPL^−/−^ flies, 18% of the brains tested showed an axonal defect (n=106). This percentage did not significantly change in the absence of both Wnt5 in APPL^−/−^ flies (21%, n=86); *p* value = 0.6027. *** indicates a *p* value< 1^-5^.

APPL and Vang are both transmembrane proteins that are part of the same receptor complex required for MB axonal growth [20]. We wondered if APPL interaction with the Wnt-PCP pathway involved a ligand and focused on *Drosophila* Wnt5 as a candidate. Wnt5 has been implicated in the regulation of MB axon growth [29,32] although the mechanism is unclear. We first examined the genetic interaction between *Wnt5* and *vang* in β-lobe axon growth. Loss of *vang* caused a highly penetrant phenotype (50%), while Wnt5 nulls showed β-lobe loss only in 5% of the brains examined, suggesting that Wnt5 is largely dispensable for β-lobe growth. Surprisingly, *Wnt5*−/−; *vang*−/− double mutants showed an almost complete rescue of *vang* loss of function (Figure 1Ca,b,d,D, Table S1). Therefore, in the absence of Vang, Wnt5 inhibits β-lobe growth, suggesting that Wnt5 interacts with another receptor and antagonizes its function in PCP-mediated axon growth. We therefore examined the genetic interaction between *Wnt5* and *appl*. Loss of Wnt5 in Appl−/− flies resulted in a phenotype similar to Appl−/− flies alone (Figure 1 Cb,c,e, E). Thus, in the absence of APPL, Wnt5 no longer negatively impacts MB axon growth, suggesting that APPL may be a Wnt5 receptor.

### APPL and human APP bind Wnt5 via the Cysteine Rich Domain

Wnt5 is a member of the large family of Wnt ligands, some of whose receptors and co-receptors harbor a conserved extracellular Cysteine Rich Domain (CRD) thought to be important for Wnt binding[33,34]. Intriguingly, APP harbors a CRD-like domain[35]in its extracellular region that includes 12 cysteine residues conserved across APP paralogs and orthologs (Figure 2A). The distribution of the cysteine residues resembles those present in the CRDs of other PCP receptors such as Fz and Ror-2 (Figure S1). We asked whether the CRDs of APP and APPL are potential Wnt5a-binding domains. To test this, we generated forms of human APP (hAPP) and APPL lacking the CRD (hAPPΔCRD and APPLΔCRD). Next, we overexpressed a tagged form of Wnt5a together with hAPP, APPL, hAPPΔCRD or APPLΔCRD in HEK293 cells and performed co-immunoprecipitation(IP) assays. Wnt5a immunoprecipitated full-length hAPP and APPL but not hAPPΔCRD or APPLΔCRD (Figure 2B). Reciprocally, full-length hAPP and APPL immunoprecipitated significant amounts of Wnt5a in contrast to hAPPΔCRD and APPLΔCRD (Figure 2B’). Similarly, APPL was found to precipitate Wnt5 from transfected *Drosophila* S2 cell lysates (Figure S2).

**Figure 2:**
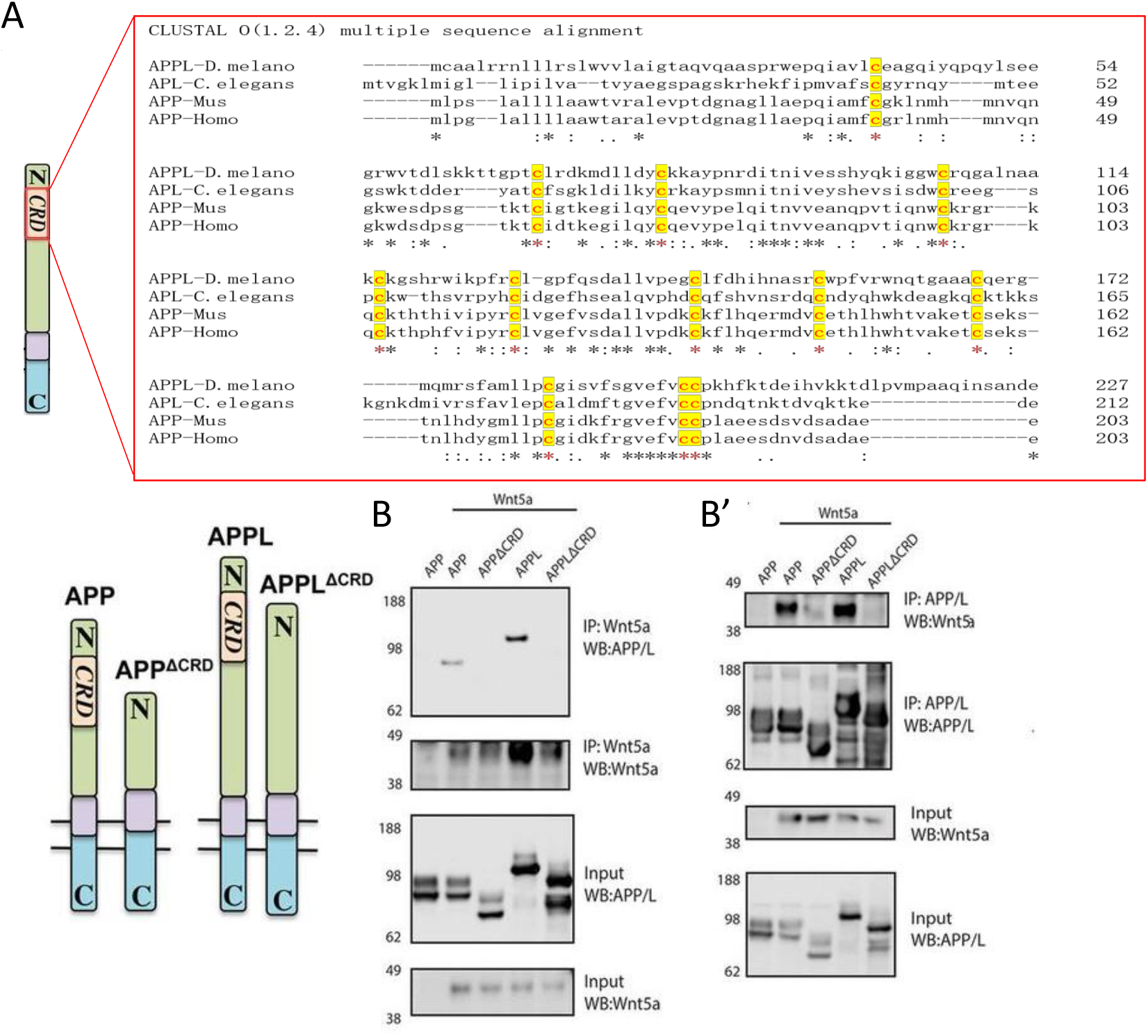
APPL and Wnt5 interact via the APP Cysteine Rich Domain. (A) APPL extracellular region contains a conserved CRD. The figure shows a CLUSTAL alignment of the CRD of different APP homologs. The 12 cysteine residues (as indicated by the red asterisks) are highly conserved across species. (B-B’) Wnt5a binds APPL and APP in a CRD dependent manner. (B) Co-immunoprecipitation (co-IP) of the full-length proteins APP-flag and APPL-flag but not their truncated forms APP^ΔCRD^-flag and APPL^ΔCRD^-flag with Wnt5a-myc. (B’) Reciprocal co-IP showing that Wnt5a-myc is co-IPed with flAPP-flag and APPL-flag can but not when the CRDs are deleted.

### Wnt5a treatment affects APP trafficking and expression in maturing mouse primary cortical neuron

The findings above suggest that the APP family may represent a new class of conserved Wnt receptors. We sought to investigate this further at a cell biological level using developing mouse embryonic primary cortical neurons as a model system. APP trafficking and processing have been intensively investigated in studies relating to AD, and according to early reports the half-life of APP is quite short, ranging from 1 hour to 4 hours [36,37]. In mouse embryonic (E16) primary neuron cultures, full-length mouse APP (fl-mAPP; henceforth we refer to mouse APP as mAPP and to human APP as hAPP) expression significantly dropped after 2 hours of treatment with translational inhibitor (Cycloheximide) (Figure S3), suggesting relatively rapid turnover of mAPP. To study the relation between mAPP and Wnts we first verified that mAPP also binds Wnt5a through its CRD and found that fl-mAPP but not mAPPΔCRD coIP’s with Wnt5a, similar to APPL and hAPP (Figure 3A). Next, we used immunofluorescence to localize mAPP with or without Wnt5a treatment in developing cortical neurons during axonal outgrowth (DIV7). mAPP is modified to maturation in the Trans Golgi Network (TGN) to be subsequently transferred to the plasma membrane where it can be internalized into early endosomes. From the early endosome, mAPP is either recycled back to the TGN through retromer-dependent sorting, or to the late endosome and then lysosome to be degraded [17,38]. We used markers for early endosomes (Rab5), TGN (Golgin97) and lysosomes (Lamp1) to trace mAPP trafficking after Wnt5a treatment. As shown in the Figure 3 (B, C), compared to control, the fraction of mAPP co-localizing with early endosomes was not affected by Wnt5a treatment, indicating normal initial internalization of mAPP. However, we found less mAPP in the TGN, and more mAPP in lysosomes (Figure 3B, D, E) suggesting that Wnt5a regulates intracellular targeting of mAPP after internalization. Importantly, the levels of expression of these markers (Rab5 Golgin97 and Lamp1) are not affected by Wnt5a treatment (Figure S4). Next, we asked if this altered trafficking affected mAPP levels. We found that the level of fl-mAPP was significantly reduced after 4hrs of Wnt5a, as shown by western blot (Figure 3F, G), with no effect on mAPP mRNA levels (Figure 3 H). The results of IF and WB suggest that the decrease of the mAPP upon Wnt5a treatment is caused by lysosomal degradation. To confirm this, we used Bafilomycin-A in combination with Wnt5a treatment to inhibit the lysosome and found that this restored mAPP to control levels (Figure 3 I-L). These data suggest that non-canonical Wnt5a-PCP signaling reduces mAPP stability.

**Figure 3:**
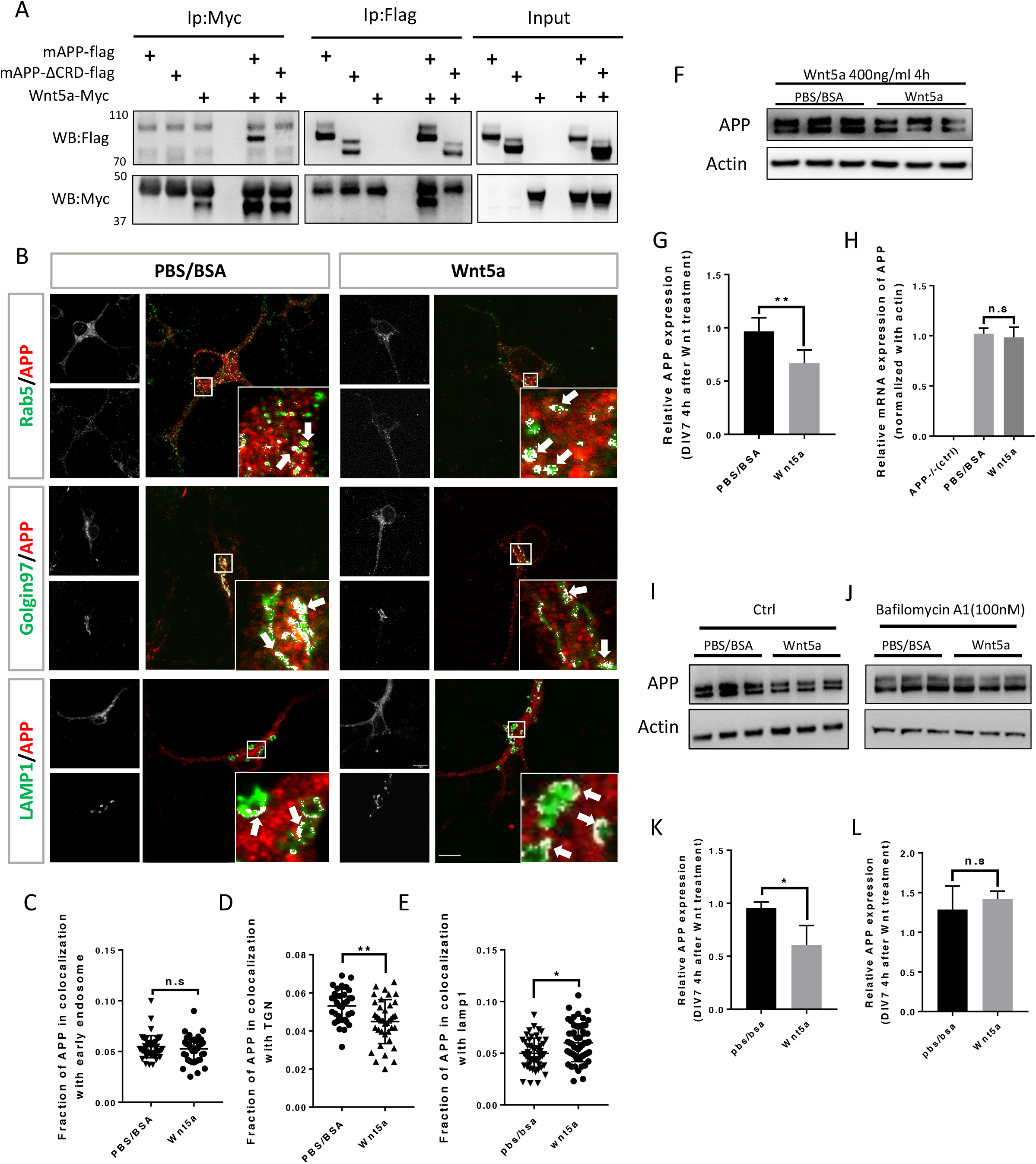
Wnt5a regulates APP expression through changing its intracellular trafficking. (A) Co-immunoprecipitation (co-IP) of Wnt5a-Myc with full-length proteins mAPP-flag or mAPP-delatCRD-flag. The tagged proteins were co-expressed in HEK293T cells and immunoprecipiteted with ant-flag or anti-Myc antibody, wild type mAPP pulled down Wnt5a and vice versa, while mAPP lacking the CRD showed impaired ability to pull down Wnt5a even with higher protein levels compared to wild type mAPP in the input. (B) mAPP localization after 4 hours PBS/BSA or Wnt5a treatment. Immunofluorescence using antibodies to APP, Rab5 (early endosome marker) Golgin97 (TGN marker) or Lamp1 (lysosome marker) to reveal mAPP localization in different intracellular compartments, the inset shows a high magnification image of the area in the white box and arrows indicate the overlap of mAPP with respective cellular compartment marker. (C-E) Quantification of the overlap between mAPP and early endosome TGN or lysosome respectively after Wnt5a treatment. (F) mAPP protein expression is altered after Wnt5a treatment, Western blotting for mAPP and Actin was done on lysates from DIV7 primary cortical neurons. (G) Quantification of the Western blot result for fig F. (H) *mAPP* mRNA is not affected after Wnt5a treatment, qPCR for *mAPP* and *actin* was done in mRNA sample from DIV7 primary cortical neurons, *APP−/−* mice derived primary neurons were used as a negative control. (I-J) The lysosome inhibitor Bafilomycin rescues Wnt5a-induced mAPP reduction in mAPP protein levels. (I) untreated controls. (J) Cells treated with the Bafilomycin. (K-L) Quantification of the western blot result for fig(I-J). Bars represent the mean±s.e.m. for at least three independent experiments. 40-50 cells from at least two independent experiments were analyzed for each group. *P<0.05, **P<0.01. Scale bar = 10um.

### Wnt3a binds to and stabilizes APP via the CRD

We wondered whether mAPP can also bind other members of the Wnt family of ligands. Wnt3a is one of the 19 Wnt members in mouse and human. During development Wnt3a usually induces β-catenin signaling pathway which plays an import role in gene expression, cell proliferation and differentiation [39,40]. Recent studies suggest that Wnt3a and beta-Catenin signaling may be involved in AD pathology [41,42]. More interestingly, studies on mouse AD models showed that Wnt3a and Wnt5a interact competitively and antagonistically with regards to APP-mediated synapse loss [23,24]. We therefore wondered whether, like Wnt5a, Wnt3a also binds to mAPP through the conserved CRD and regulates its levels. To test this, we performed IP experiments with Wnt3a. We found that fl-mAPP and Wnt3a co-IP in a CRD-dependent fashion (Figure 4A). We next tested the effects of Wnt3a treatment on APP trafficking. As shown in Figure (4 B, C), the fraction of mAPP colocalized with early endosomes was not affected. However, more mAPP was present in the TGN compared to controls (Figure 4 B, D), with no effect on the lysosomal mAPP fraction (Figure 4 B, E). The expression levels of Rab5, Golgin97 and Lamp1 are not affected after Wnt3a treatment (Figure S4). Western blot analysis showed increased fl-mAPP upon Wnt3a treatment (Figure 4 F, G), but no effect on mRNA levels (Figure 4 H). There is evidence that mAPP is recycled back to the TGN from early endosomes through the retrograde pathway[38]. To test whether Wnt3a regulates mAPP retrotrafficking to the TGN, we co-treated primary neurons with Wnt3a and a retromer inhibitor (LY294002). This reversed the effect of Wnt3a on mAPP trafficking protein levels (Figure 4 I-L). Finally, we tested the effects of simultaneous treatment with Wnt3a and Wnt5a. This resulted in no change to APP protein levels compared to controls, suggesting that Wnt3a and Wnt5a neutralize each other’s effects on mAPP (Figure 4 M, N), again with no effects on mRNA levels (Figure 4 O). Taken together, these data indicate that Wnt3a also binds to mAPP via the CRD and regulates mAPP trafficking and expression and that Wnt5a and Wnt3a act antagonistically to regulate APP protein homeostasis.

**Figure 4:**
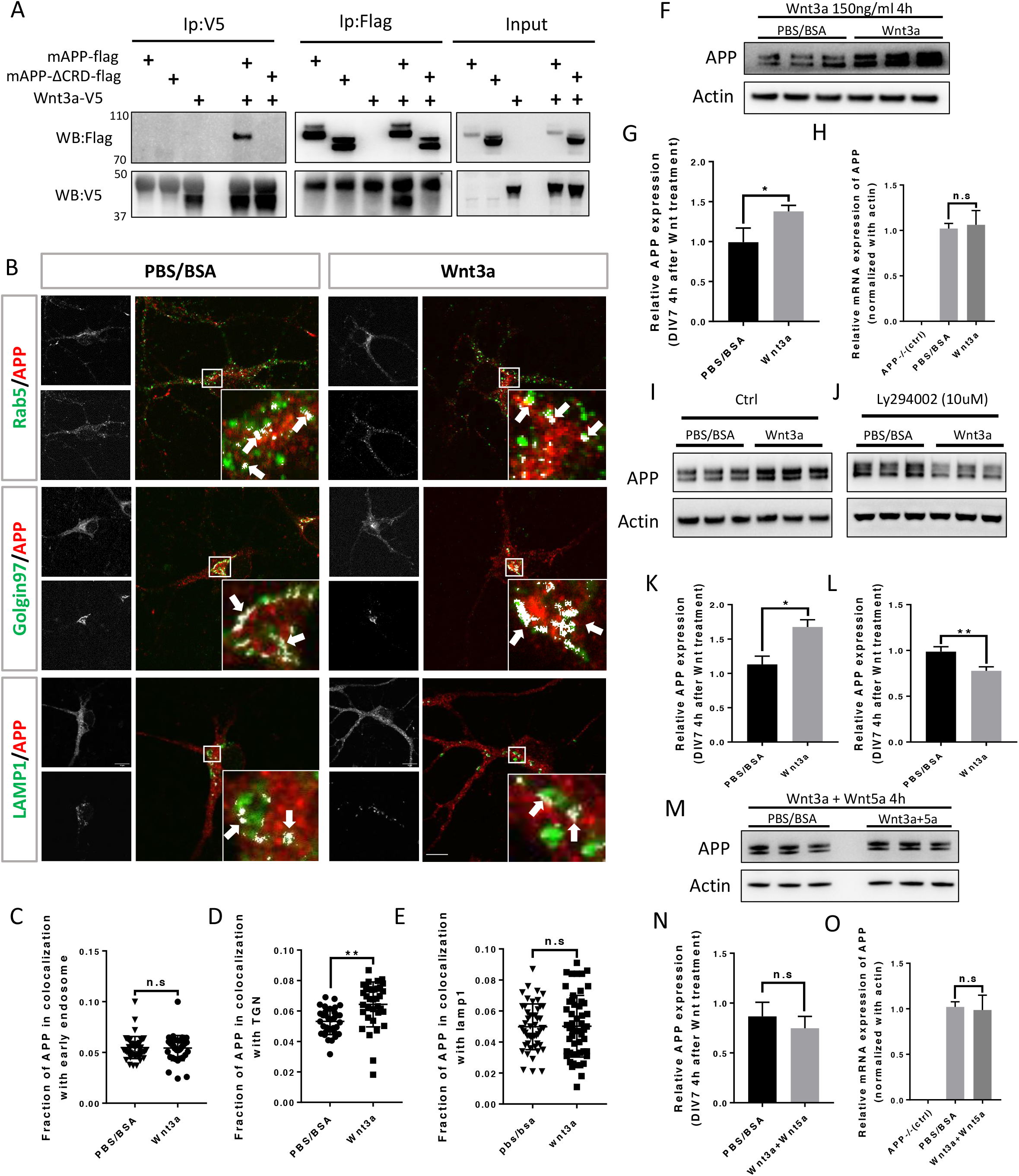
Wnt3a binds to and regulates APP expression through changing its intracellular trafficking. (A) Co-immunoprecipitation (co-IP) of Wnt3a-V5 with full-length proteins mAPP-flag or mAPPL ΔCRD. The tagged proteins were co-expressed in HEK293T cells and immunoprecipiteted with ant-flag or anti-v5 antibody, wild type mAPP pulled down Wnt3a and vice versa, while mAPP lacking the CRD showed impaired ability to pull down Wnt3a even with higher protein levels compared to wild type mAPP in the input. (B) mAPP localization after 4 hours PBS/BSA or Wnt3a treatment. Immunofluorescence using antibodies to APP, Rab5, Golgin97 or Lamp1 to reveal mAPP localization in different intracellular compartments, the inset shows a high magnification image of the area in the white box and arrows indicate the overlap of mAPP with respective cellular compartment marker. (C-E) Quantification of the overlap between mAPP and early endosome, TGN or lysosome, respectively, after Wnt3a treatment. (F) mAPP protein expression after Wnt3a treatment. Western blotting for mAPP and Actin was done on lysates from DIV7 primary cortical neurons. (G) Quantification of the Western blot results for fig F. (H) *mAPP* mRNA is not affected after Wnt3a treatment, qPCR for *mAPP* and *actin* was done in mRNA sample from DIV7 primary cortical neurons, *APP−/−* mice derived primary neurons were used as a negative control. (I-J) The Retromer inhibitor Ly294002 rescues Wnt3a-induced increase in mAPP expression levels (I) untreated controls. (J) Cells treated with Ly294002. (K-L) Quantification of the Western blot result for fig(I-J). (M-O) Wnt5a and Wnt3a working in a competing way on affecting mAPP protein expression. (M) Western blot performed with cell lysate from the DIV7 primary cortical neuron treated with Wnt3a and Wnt5a at the same time for 4 hours, PBS/BSA group act as control group. (N) Quantification of the Western blot result for fig M. (O) qPCR results of mAPP knockout neurons (negative control) and PBS/BSA or Wnt3a+Wnt5a treated neurons. Bars represent the mean±s.e.m. for at least three independent experiments. 40-50 cells from at least two independent experiments were analyzed for each group. *P<0.05, **P<0.01. Scale bar = 10um.

### The CRD is required for Wnt-mediated regulation of APP trafficking and expression

Our data thus far show that APP interacts with Wnts through its CRD and that Wnts regulate APP intracellular trafficking and expression. We therefore asked whether the CRD is required for the effects of Wnts on mAPP. To address this question, we created two lentiviral vectors: pLv-pSyn1-mAPP-Flag-IRESeGFP (flag-tagged fl-mAPP) and pLv-pSyn1-mAPPΔCRD-Flag-IRESeGFP (flag-tagged mAPPΔCRD). Primary cortical neurons from APP knockout mice were transduced with the fl-mAPP or mAPPΔCRD vectors, or a control GFP vector (pLv-pSyn1-IRESeGFP) exogenous APP/APPΔCRD could be well detected using either anti-APP or anti-flag antibodies (Figure S5 A-C). Virus induced APP/APPΔCRD expression level was on average slightly lower than endogenous (Figure S5 B, C) which is largely explained by ~50% transduction efficiency (data not shown). In neurons transduced with wild type mAPP we confirmed that mAPP expression could be increased and decreased by Wnt3a and Wnt5a treatments, respectively (Figure 5 A,C). In contrast, in neurons transduced with mAPPΔCRD those effects were eliminated (Figure 5 B,D). Finally, we performed IF to trace mAPP and mAPPΔCRD localization. Neurons transduced with wild type mAPP showed the same results as wild type neurons with more mAPP in the TGN upon Wnt3a treatment and more mAPP in lysosomes upon Wnt5a treatment (Figure S6 A-D). In contrast, neurons transduced with mAPPΔCRD neither Wnt3a, nor Wnt5a treatment showed a significant effect on mAPP localization to early endosomes, TGN or lysosomes compared to controls (Figure 5 E-H). In summary, these data show that the CRD of mAPP is critical for Wnt3a/5a binding and mediates the effects of Wnts on mAPP trafficking and expression.

**Figure 5:**
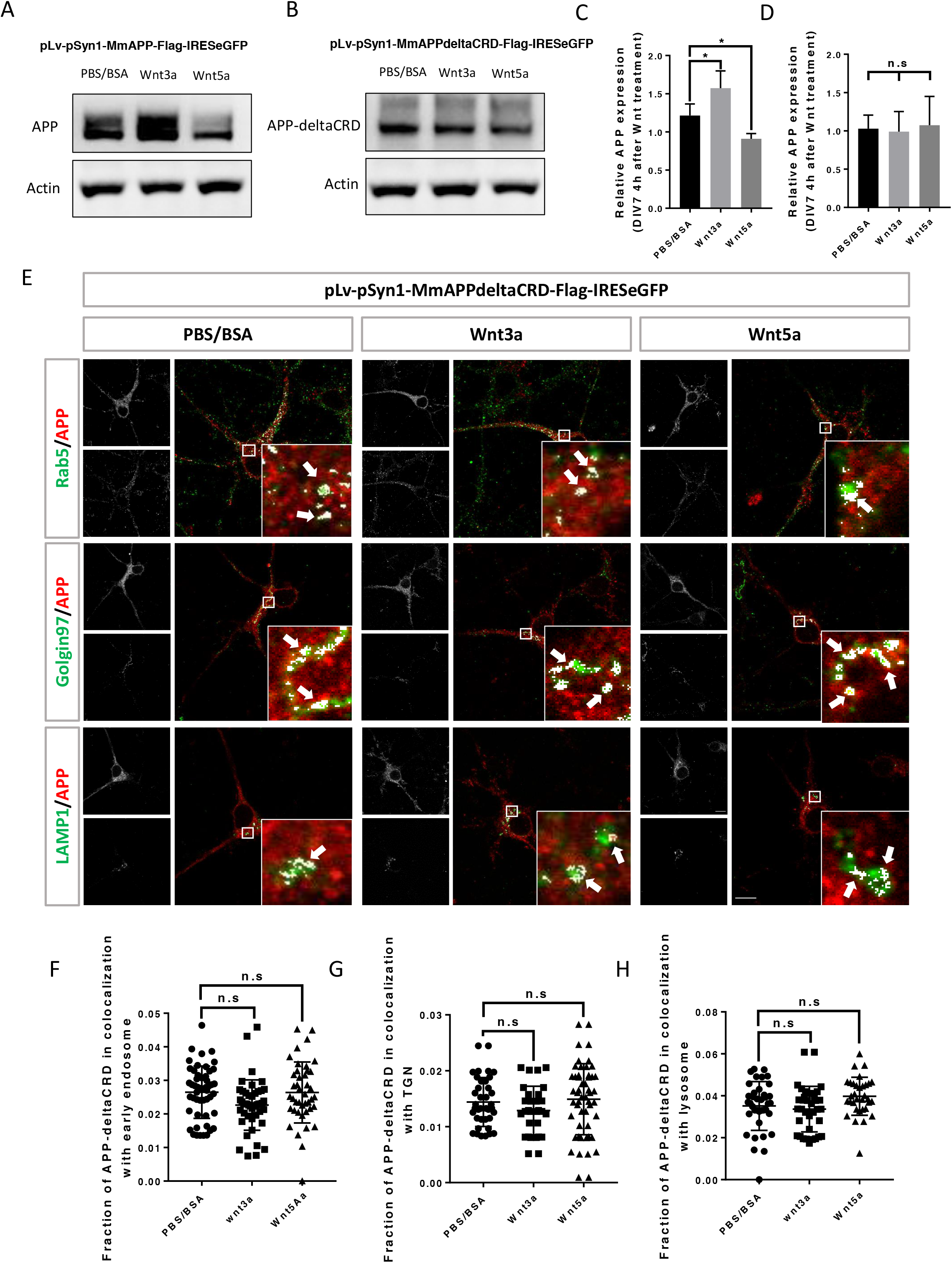
CRD is required for Wnt3a/5a to affect APP trafficking and expression. (A-D) CRD is critical for Wnt3a/5a regulation of mAPP protein expression. (A-B) APP knock-out primary cortical neuron expressing exogenous wild type or CRD mutant mouse APP via lenti-virus transduction were treated with Wnt3a or Wnt5a at DIV7, 4 hours later protein samples were collected for Western blots. Wnt3a upregulated fl-mAPP and Wnt5a downregulated mAPP (A), in contrast both Wnt3a or Wnt5a failed to affect mAPPΔCRD expression (B). (C-D) quantification of the Western blots results for figure A and B respectively. (E-H) Routing of mAPP trafficking by Wnt3a/5a requires the CRD. (E) Localization of exogenous mAPP in APP knock-out primary cortical neurons after 4 hours of Wnt3a or Wnt5a treatment. Immunofluorescence using antibodies to APP, Rab5 (early endosome marker) Golgin97 (TGN marker) or Lamp1 (lysosome marker) to reveal mAPP localization in different intracellular compartments, the inset shows a high magnification image of the area in the white box and arrows indicate the overlap of mAPP with respective cellular compartment marker. (F-H) Quantification of the overlap between mAPP and early endosome, TGN, or lysosome, respectively, after Wnt3a or Wnt5a treatment. Bars represent the mean±s.e.m. for at least three independent experiments. 40-50 cells from at least two independent experiments were analyzed for each group. *P<0.05, Scale bar = 10um.

### CRD is critical for APP to regulate neurite outgrowth and complexity

APP and its proteolytic products have been reported to affect neurite outgrowth during development [43,44] in different systems. We used primary cortical neuron derived from E16.5 mice embryos to investigate if the CRD of mAPP is required for regulation of neurite outgrowth by mAPP. We examined axonal and dendritic outgrowth (Figure 6A) at three developmental stages *in vitro:* DIV2, DVI3 and DIV7[45].

**Figure 6:**
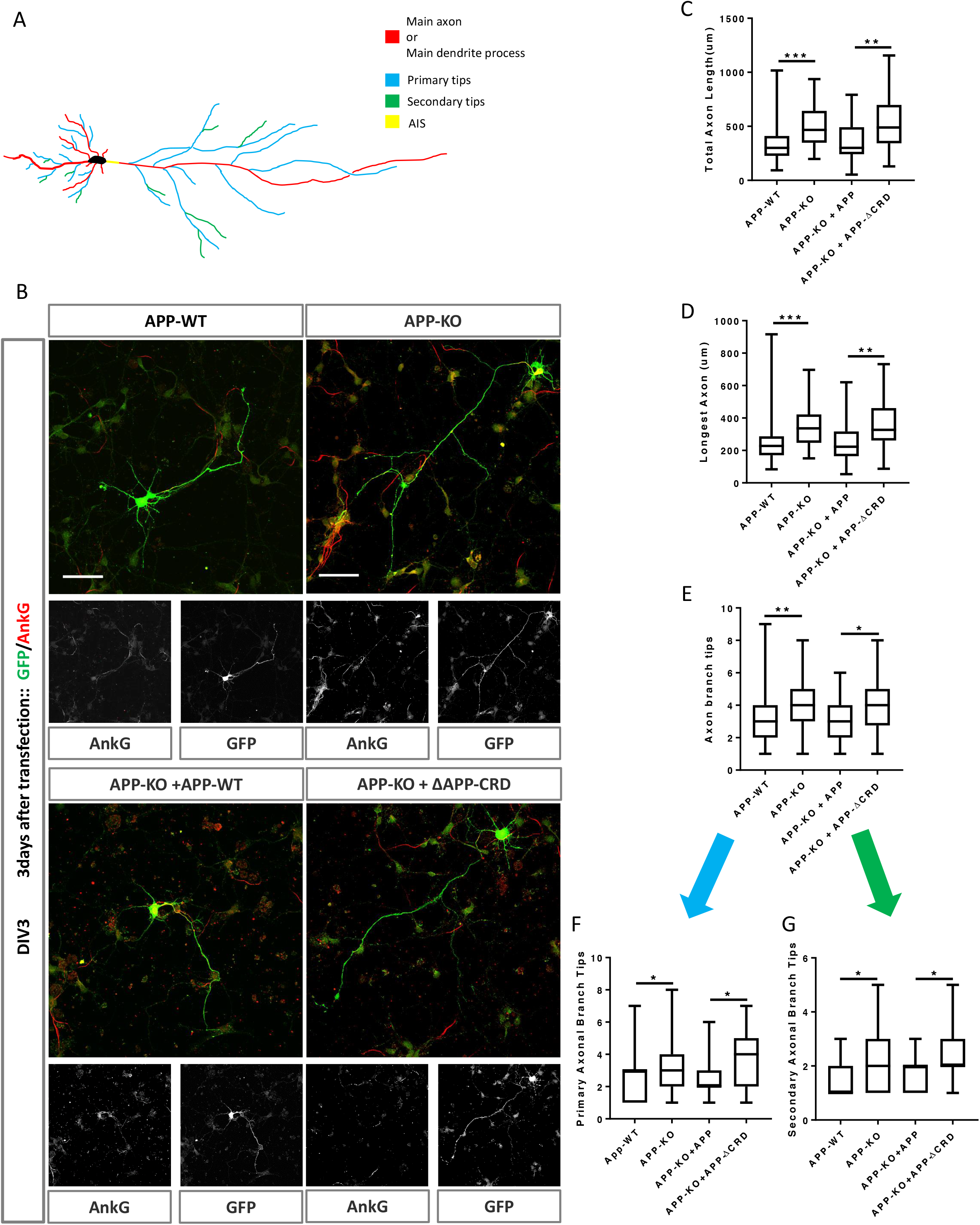
Cysteine Rich Domain is critical for APP to regulate neurite outgrowth at DIV3. (A) Schematic of a primary neuron, colored lines indicate axonal or dendritic branch tips which were quantified, yellow indicates the Axon Initial Segment (AIS) marked with Ankry G in experiments (B) Representative confocal images primary cortical neurons of the four genotypes examined: mAPP wild type, mAPP knock out and mAPP knock-out rescue with APP or CRD-mutant APP. Transfection of plasmid containing GFP alone, mAPP-flag-GFP or mAPPΔCRD-flag-GFP performed at the onset of cell seeding. Cells were fixed at DIV3 and immunolabeled with GFP, AnkG. (C-E). Quantification of three parameters from the four primary neuron genotypes at DIV3. (C) Quantification of the total axon length (the main axonal process and the branches deriving from the main process) at DIV3 (D) Quantification of the length of longest axonal process at DIV3. (E) Quantification of the total axonal branch tips at DIV3. (F) Quantification of primary branch tips at DIV3. (G) Quantification of secondary branch tips at DIV3. 50-60 cells from at least two independent experiments were analyzed for each group. *P<0.05, **P<0.01 ***P<0.001, Scale bar = 50um.

While we found no effect on initial outgrowth at DIV2 (Figure S7A-F), at DIV3 mAPP knockout neurons exhibit increased axonal outgrowth compared to controls reflected in three parameters: total axon length, longest axon length and the number of branch tips (Figure 6B-E). In contrast, dendritic outgrowth was not different from controls (Figure S8A-C). We asked whether the CRD was required for mAPP function during neurite outgrowth. To this end we transfected mAPP knockout neurons with either fl-mAPP or mAPPΔCRD. Increased axonal length and axonal branch tips were rescued by the fl-mAPP but not by the form lacking the CRD at DIV3 (Figure 6B-E). Next, we analysed axonal branching in greater detail and found that loss of mAPP increased the numbers of both primary and secondary axonal branches at DIV3, an increase that was rescued by fl-mAPP but not by mAPPΔCRD (Figure 6F,G). Finally, we examined the Axon Complexity Index (ACI) [46], which measures the ratio of branches of different orders to total branch number, at DIV3. At this early stage, the ACI showed a tendency to increase in mAPP knockout neurons that was not significant (Figure S8D), likely because both primary and secondary branches show a similar level of increase. Together these data suggest an overall increase in axonal growth. In contrast to axonal growth, we found no significant alterations in dendritic length or branching (Figure S8E-G) consistent with the fact that the spur in dendritic outgrowth is largely initiated at DIV4[47,48].

To further analyse neurite outgrowth, we examined axonal and dendritic growth at DIV7. By this stage mAPP knockout neurons showed an increased ACI (Figure 7A,B). In contrast, total axonal length, longest axon length and the total number of branch tips was not significantly different (Figure 7C-E). The increase in axonal complexity in mAPP knockout neurons was due to a significant reduction in the number of primary branches and a significant increase in the number of secondary branches (Figure 7F,G). Once again, all phenotypes were rescued by fl-mAPP but not mAPPΔCRD.

**Figure 7:**
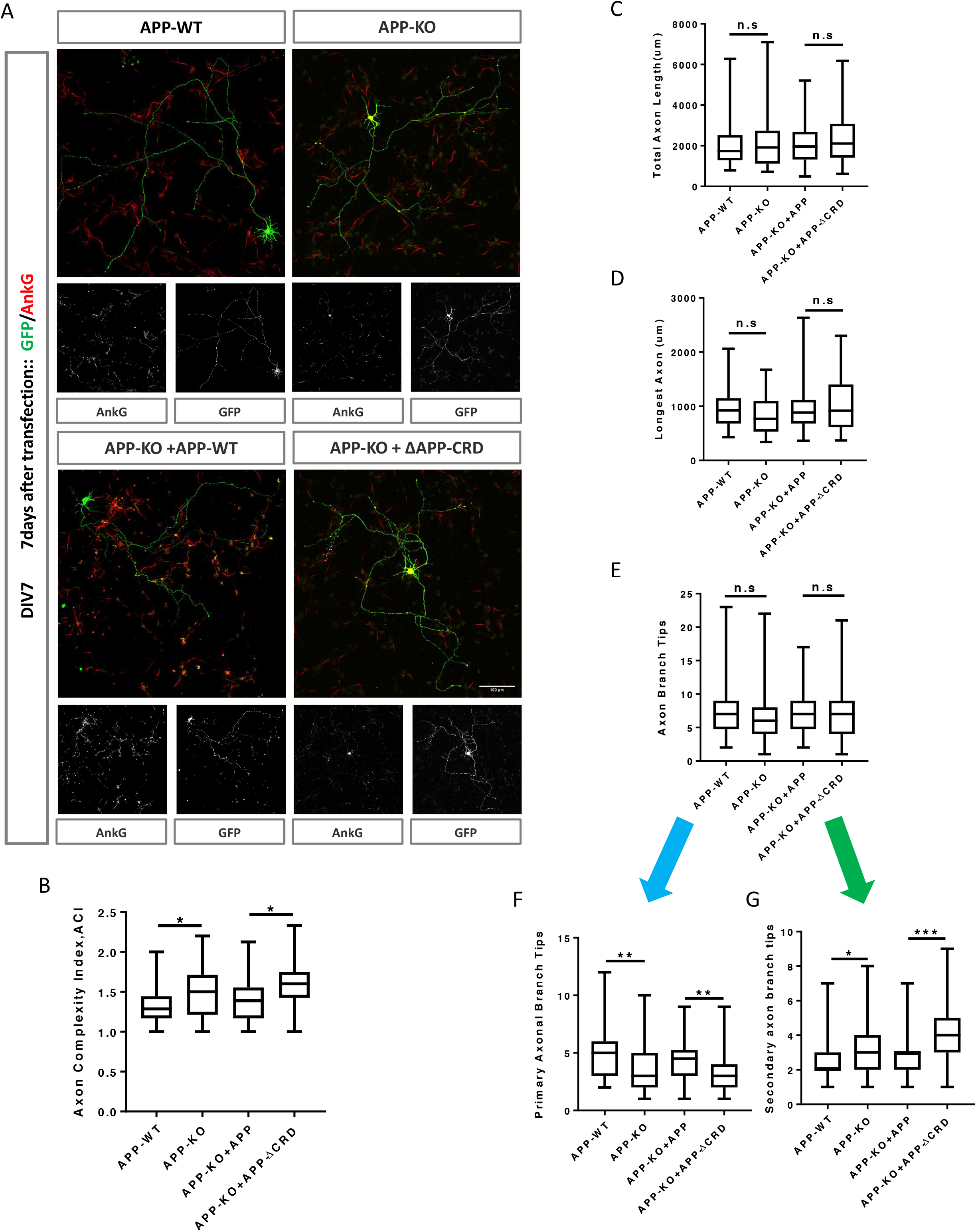
Cysteine Rich Domain is critical for APP to regulate neurite outgrowth and complexity at DIV7. CRD is required for APP to regulate axon branching complexity at DIV7. (A) Representative confocal images showing primary cortical neurons of the four genotypes examined: mAPP wild type, mAPP knock out and mAPP knock-out rescued with APP or CRD-mutant APP. Transfection of plasmid containing GFP alone, mAPP-flag-GFP or mAPPΔCRD-flag-GFP performed at the onset of cell seeding. Cells were fixed at DIV7 and immunolabeled with GFP, AnkG. (B) Analysis of Axon complexity Index (ACI) at DIV7. (C) Quantification of the total axonal length at DIV7. (D) Quantification of the length of longest axonal process at DIV7. (E) Quantification of the all axonal branch tips at DIV7. (F) Quantification of primary branch tips at DIV7. (G) Quantification of secondary branch tips at DIV7. 50-60 cells from at least two independent experiments were analyzed for each group. *P<0.05, **P<0.01 ***P<0.001, Scale bar = 100um

Finally, we examined dendritic growth at DIV7. We observed no difference in total dendrite length or the size of the longest dendrite (Figure S9A,B), but observed a significant decrease in the total number of dendritic processes in mAPP knockout neurons compared to controls (Figure S9C). This reduction was due to the presence of fewer main dendritic processes in mAPP knockout neurons but no effect was observed on the primary or secondary dendritic branches (Figure S9D-F). All phenotypes were rescued by fl-mAPP but not mAPPΔCRD. Taken together, our results show that the role of APP in neuronal maturation requires the CRD domain.

## DISCUSSION

Here we identify a previously-unknown conserved Wnt receptor function for APP proteins. We show that APP binds both canonical and non-canonical Wnt ligands via a conserved cysteine rich domain and that this binding regulates the levels of full length APP by regulating its intracellular trafficking from early endosomes to the trans Golgi network versus the lysosome. Finally, we show that APP through the CRD regulates neurite growth and axon branching complexity in primary mouse cortical neurons

APP has been extensively reported to be involved in regulating neurite outgrowth [43,44,49–52], with conflicting conclusions as to whether APP promotes or inhibits neurite outgrowth. In our experiments, we found that while in *Drosophila* APPL loss reduced axonal growth, the comparison of axonal outgrowth and branching in primary cortical neuron derived from mAPP wild type or mAPP knock out mice at DIV2, DIV3 and DIV7 showed that loss of mAPP significantly accelerated axonal maturation. Specifically, we found that the initial phase of axonal growth at DIV2 is unaffected, but that APP mutant axons grow longer at DIV3 and then show increased axon complexity at DIV7. We therefore suggest that the conflicting data in the literature may arise from examining different types of neurons at different time points, where the requirement of APP may differ in a context-specific manner. We speculate that this context specificity may in part be due to the levels and types of Wnt ligands present in the environment.

Finally, our findings suggest that in addition to the well described proteolytic processing of APP, the regulation of its recycling by Wnt ligands may be crucial for its function. It is important to note that, like proteolytic processing, Wnt ligands regulate APP stability post-translationally, as we found no effect on APP mRNA levels upon Wnt treatment. With regards to the role of APP processing in Alzheimer’s disease, recently published work suggests that an imbalance between Wnt3a/canonical signaling pathway and the Wnt5a/PCP signaling pathway at the initial step of amyloid beta production could trigger a vicious cycle favouring the amyloidogenic processing of APP [23,24]. We suggest that our data provide a mechanistic framework for understanding how this may occur in neurons.

## ACKNLOWEDGMENTS

We thank Dr. Radoslaw Ejsmont for writing the co-localization macro, Natalia Danda for the construction of the plasmids used in this work, Drs. Ariane Ramaekers, Natalia Mora Garcia, Gerit Linneweber, Simon Weinberger and Guangda Liu for helpful discussions. We thank Drs. Zeynep Kalender Atak and Marina Naval Sanchez for support on the statistical analysis of the data and Dr. Jean-Maurice Dura for fly lines. Mouse breeding work was conducted at the PHENO-ICMice facility. The Core is supported by 2 Investissements d’Avenir grants (ANR-10-IAIHU-06 and ANR-11-INBS-0011-NeurATRIS) and the “Fondation pour la Recherche Médicale”. Primary neuron culture work was carried out at the CELIS core facility with support from Program Investissements d’Avenir (ANR-10-IAIHU-06). Light microscopy was carried out at the ICM.Quant facility. We thank all core technical staff involved. This work was supported by ICM, the program “Investissements d’avenir” ANR-10-IAIHU-06, the Einstein-BIH program, the Paul G. Allen Frontiers Group, and the Roger De Spoelberch Foundation (BAH), the Vlaams Instituut voor Biotechnologie (VIB; BAH and BDS), the Methusalem grants from KU Leuven (BDS and BAH), Fonds Wetenschappelijke Onderzoeks (FWO) grants G.0543.08, G.0680.10, G.0681.10 and G.0503.12 (BAH), the Nederlandse Organisatie voor Wetenschappelijk Onderzoek (NWO; ZonMw TOP grant 40-00812-98-10058) and the Hersenstichting Nederland [HS 2011(1)-46] (LGF), grant “projet ARC n° SFI20121205950” from the Association pour la Recherche sur le Cancer (ARC, JMD) and a doctoral fellowship from the Centre National de Recherche Scientifique Libanais (LCNRS, MN). Tengyuan Liu and Tingting Zhang are funded by the Chinese Scholarship Council (CSC). The authors declare no competing financial interests.

## AUTHOR CONTRIBUTIONS

The authors have made the following declarations about their contributions: Conceived and designed the experiments: Tengyuan Liu, Tingting Zhang, Maya Nicolas, Lee G. Fradkin and Bassem A. Hassan. Performed the experiments: Tengyuan Liu, Tingting Zhang, Maya Nicolas, Heather Rice, Alessia Soldano, Annelies Claeys, Iveta M. Petrova and Jean-Maurice Dura. Analyzed the data: Tengyuan Liu, Tingting Zhang, Maya Nicolas, Heather Rice, Bart De Strooper, Lee G. Fradkin, and Bassem A. Hassan. Wrote the paper: Tengyuan Liu, Maya Nicolas, and Bassem A. Hassan. All authors read and approved the manuscript

## Materials and Methods

### Drosophila stocks and maintenance

Flies were raised at 25°C, on standard cornmeal and molasses medium. The stocks used in this study are: w*, Appl^d^; Vang^stbm-6^; w1118, Wnt5^400^; P247Gal4; w*, Appl^d^,Wnt5^400^.

### Cloning

All constructs were generated by PCR amplification and overlap extension PCR. PCR products were inserted into the respective vectors by classical restriction enzyme cloning. All constructs were sequence-verified. To generate transgenic flies, open reading frames with epitope tags were cloned into the pUAST-attB fly expression vector and transgenes were inserted into the genome at the VK37 docking site (2L, 22A3) via PhiC31-mediated transgenesis.

### Mushroom body analyses

Adult fly brains were dissected in phosphate buffered saline (PBS) and fixed in 3.7% formaldehyde in PBT (PBS+ 0.1% Triton100-X) for 15 min. Then, the brains were washed 3 times in PBT and blocked in PAX-DG for 1 hr at RT. The samples were later incubated with the primary antibody overnight at 4°C. After incubation, the brains were washed 3 times with PBT and incubated with an ALEXA Fluor^®^ secondary antibodies (Life technologies) for 2 hr at RT. After 3 times washes in PBT, the brains were mounted in Vectashield (Vector Labs, USA) mounting medium. The following antibodies were used: mouse anti-FasII (Developmental Studies Hybridoma Bank (DSHB), 1/50), rabbit anti-GFP (Invitrogen, 1/500), rat anti-Cadherin (DSHB, 1/100). The mounted brains were imaged on a LEICA DM 6000 CS microscope coupled to a LEICA CTR 6500 confocal system and a Nikon A1-R confocal (Nikon) mounted on a Nikon Ti-2000 inverted microscope (Nikon). The pictures were then processed using ImageJ

### Primary cortical neuron culture, virus transduction and plasmids transfection

The animal experiments were carried out in accordance with animal welfare regulations and have been approved by Ethic Committee and French regulatory authorities of the respective institutes. APP knock out mice were a gift from the De Strooper lab. Cortical primary neuron cultures were prepared from embryonic day 16.5 mice (APP wild or APP mutant), as described previously[53].

Virus (pLv-pSyn1-mAPP-Flag-IRESeGFP, pLv-pSyn1-mAPP Δ CRD-Flag-IRESeGFP or pLv-pSyn1-eGFP) transduction was performed during seeding in 24 well plates(4×10^5^cells/mL). 50uL (50uL par well) of the adequate lentiviral dilution in the medium of interest must be ready in tubes. Seed 150uL of the cell preparation to each well. Immediately add 50ul of the diluted lentiviral preparation to each well(final MOI 2). Mix slowly the cells-lentivirus suspension by pipetting. Incubate 1h at 37°C. Finally add 800uL of culture medium to each well and incubate for 3 additional days before any analysis.

Plasmids (pLv794_pTrip_PromSynaptin1_GFP_DeltaU3, pLv-pSyn1-MmApp-FLAG-IRES-eGFP or pLv-pSyn1-mAPP Δ CRD-FLAG-IRES-eGFP) transfection was performed at the onset of cell seeding (4×10^5^cells/mL) in 24 wells plates with coverslip coated with PDL 24hour before. All procedure follow the protocol from Lipofectamine 3000 transfection reagent (Thermofisher Catalog Number: L3000008) with little modified, each well transfection with 500ng corresponding plasmid, medium was refreshed 5-6 hours after transfection.

### Wnt and inhibitor treatment in primary neuron

Wnt5a(400ng/ml)(645-WN-010, R&D Systems), Wnt3a(150ng/ml)(1324-WNP-010, R&D Systems) and PBS/BSA(control) addition performed at Div 7. In all experiments related to inhibitor, cells will be treated with inhibitor 1hour after Wnt addition(Bafilomycin A1(100nM, invivogen, 88899-55-2), LY294002(10uM, Sigma, L9908)-as wnt3a or wnt5a treatment could affect APP protein expression clearly 2 hours later (Figure S10)- and a DMSO (0.05%DMSO in culture medium) group will be set as control. Protein or RNA samples collected after 4hours Wnt treatment.

### Quantitative real-time PCR (qRT-PCR)

Cells were lysed for RNA or protein extraction and then subjected to qRT-PCR or western blot as previously described [54]. The detailed sequence of each primer used in the whole study for qRT-PCR was summarized below:β-actin, sense 5’-TCCATCATGAAGTGTGACGT-3’ and antisense 5’-GAGCAATGATCTTGATCTTCAT-3’, mAPP, sense 5’-CATCCAGAACTGGTGCAAGCG-3’ and anti-sense 5’-GACGGTGTGCCAGTGAAGATG-3’ GAPDH, sense 5’-GCTGCCAAGGCTGTGGGCAAG-3’ and anti-sense 5’-GCCTGCTTCACCACCTTC-3’.

### Western Blots

Western blots was performed follow the user guide of Mini Gel Tank (ThermoFisher, A25977) with little modified. Briefly, Protein samples collected from total cell lysates with RIPA buffer, supernatant were collected after centrifugation, denatured samples were loaded separated on the 4-12% polyacrylamide gels (SDS-PAGE)(ThermoFisher, NW04122BOX) and then transfered to the 0.42um nitrocellulose membranes, blots visualization performed after primary and secondary antibody incubation.

### Immunoprecipatation

HEK293 cells in 10cm dish (70% confluent) were transfected with pCDNA3-MmApp-FLAG-IRES-eGFP, pCDNA3-mAPP Δ CRD-FLAG-IRES-eGFP, pCDNA3-Wnt5a-myc, pCDNA-Wnt3A-V5 or co-transfected APP or APP Δ CRD with Wnt3a or Wnt5a. 3 days after transfection, cells were collected with NP-40 lysis buffer, then sample supernatant was collected after >12000 rpm centrifugation for 20min at 4 degree, 450ul supernatant was incubated with primary antibody overnight at 4 degree, then Protein G sepharose beads(Thermo Fisher Scientific) were added to the sample to capture protein-antibody complex by rotating 2 hours at room temperature, then washed four times with the lysis buffer, and resuspended with loading buffer then denatured at 95 degree for 10 mins, blots visualized after western blot procedure as described before.

### Immunofluorescence

At DIV 7, cultured primary neurons in 24 wells were washed once with 1X PBS, then fixed in 4% paraformaldehyde (PFA) in PBS at room temperature (RT) for 10 minutes. After 3 times washing with 1X PBS, cells were blocked with 10% normal donkey or goat serum in 1 X PBS for 30 minutes at RT followed by 3 times washing in 1 X PBS. Thereafter, cells were incubated with primary antibodies diluted in 1 X PBS containing 1% normal donkey or goat serum for 2-3 hours at RT. 3 times washing with 1 X PBS, incubated with appropriate secondary antibodies conjugated with Alexa Fluor 488, Alexa Fluor 555, or Alexa Fluor 647 (1:500, Invitrogen) in 1 X PBS containing 1% normal donkey or goat serum for 1 hours at RT. Washed with 1 X PBS for 3 times, then counterstained the slides with DAPI (1:2000, Sigma) and mounted by using Vectashield (Vector) after rinsing. Primary antibodies used in this study were rabbit anti-APP (1:100, Synaptic Systems, 127 003), mouse anti-rab5 (1:100, Synaptic Systems, 108011), mouse anti-Golgin-97 (1:100, Invitrogen, A-21270), rat anti-Lamp1 (1:20, Santa Cruz, sc-19992). After staining, images were obtained by using confocal microscope (Olympus FV-1200 or Leica SP8). The percentage of APP or APP-ΔCRD co-localizing with rab5, Golgin-97 and Lamp1 was calculated using JACOP [55] via an automated macro that was written in house.

### Statistical analyses

Statistical analyses were performed using GraphPad Prism software (GraphPad Software Inc., La Jolla, CA, USA). Differences between groups were compared using the G-test, One-way ANOVA and Mann Whitney two-sample test (two-tail) as appropriate.

## Supplementary-Figures

### Summary of Supplemental Material

**Table S1:**
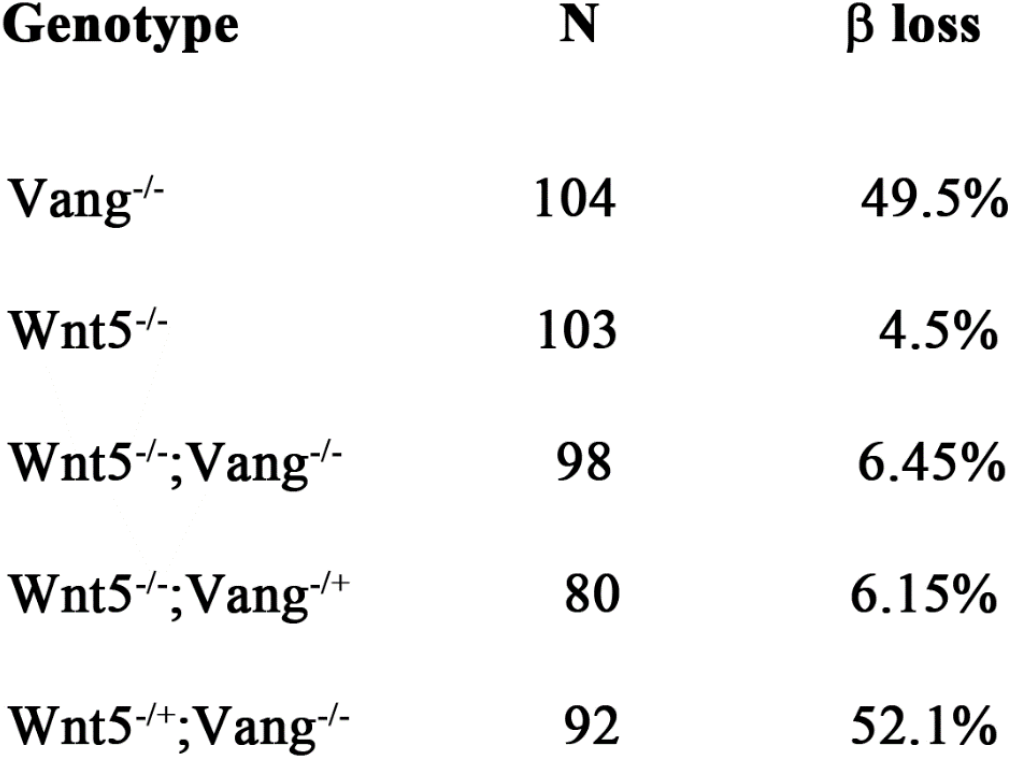
Genetic interaction between Wnt5 and Vang. The table lists the penetrance of the phenotype and the number of brains analyzed in the Vang-Wnt5 genetic interaction experiment.

**Figure S1:**
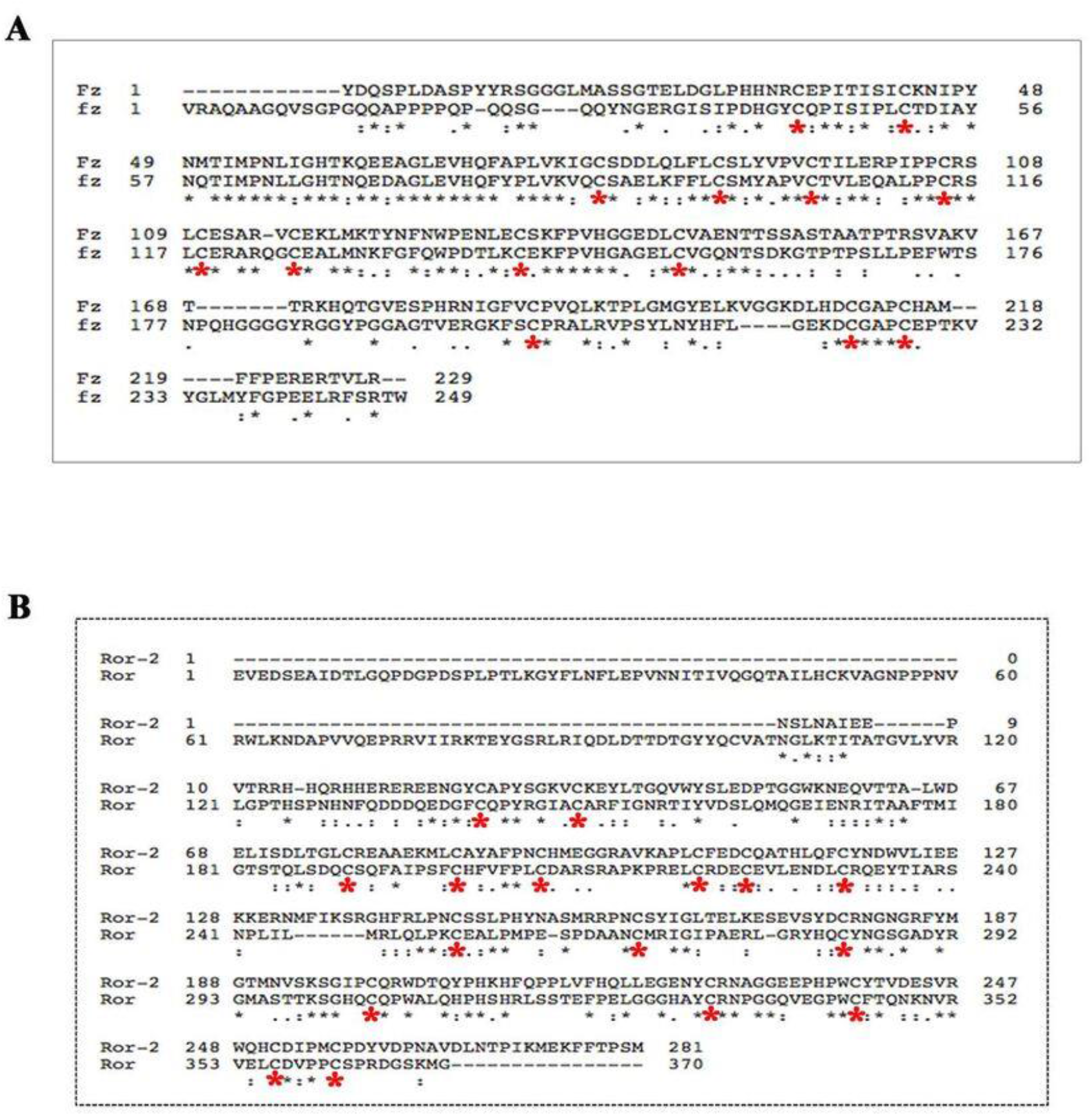
PCP receptors harbor conserved Cysteine Rich Domains (CRD) CLUSTAL alignment of the extracellular regions of Drosophila Frizzled (Fz) and Mus musculus Firzzled-1 (Fz-1) (A). CLUSTAL alignment of the extracellular regions of Drosophila Ror-2 and Mus musculus Ror-2 (Ror)(B). All proteins showed conserved cysteine residues in their extracellular region (as indicated by the red asterisks).

**Figure S2:**
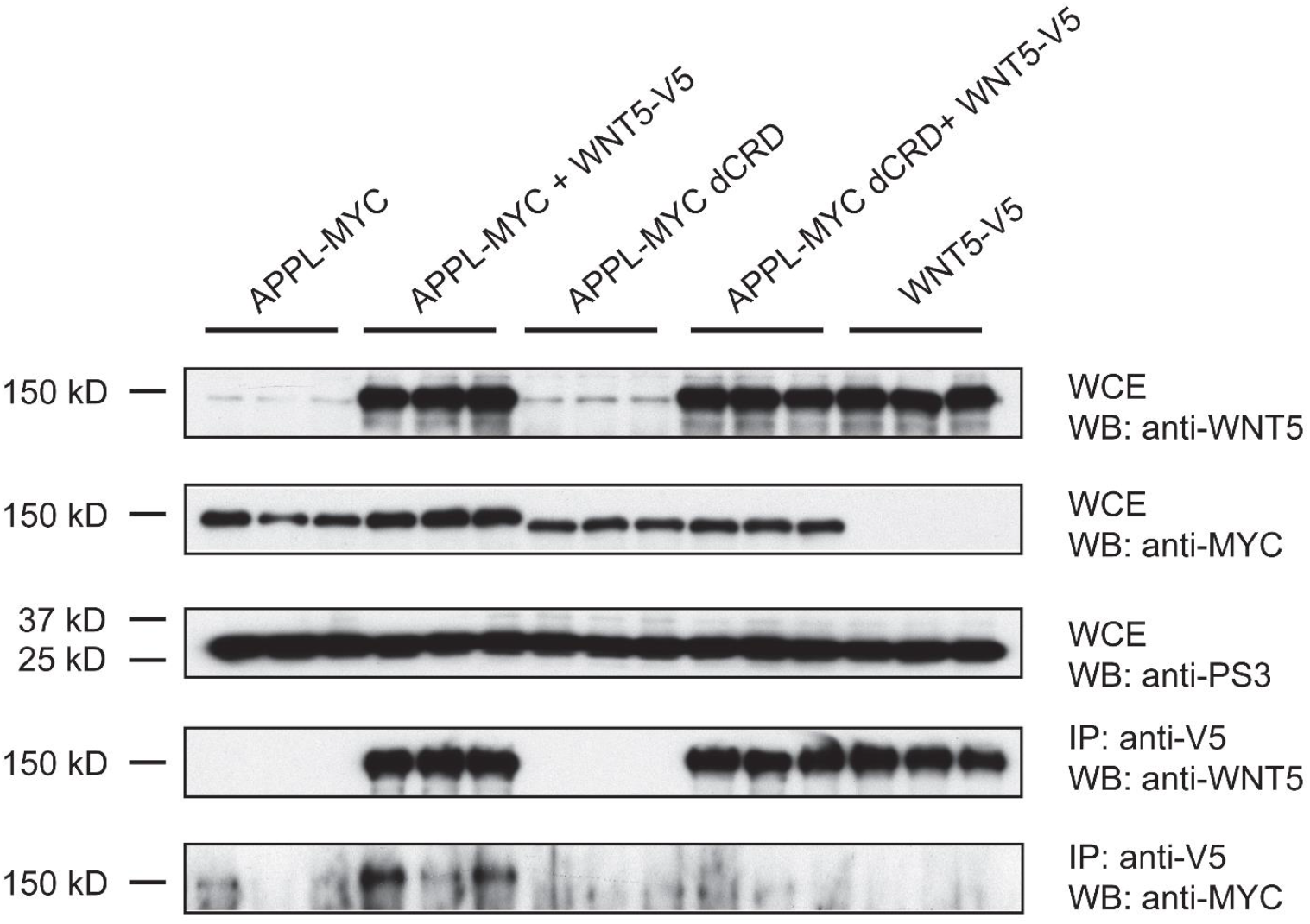
Co-immunoprecipitation assays reveal that Drosophila APPL binds to WNT5. S2 cells were transfected in triplo with the indicated expression plasmids, lysates prepared and V5-tagged WNT5-containing complexes were immunoprecipitated with anti-V5 antisera. Following SDS-PAGE and transfer to PVDF membrane, MYC-tagged APPL species were detected with anti-MY C and an HRP-conjugated chemiluminscent detection reagent

**Figure S3:**
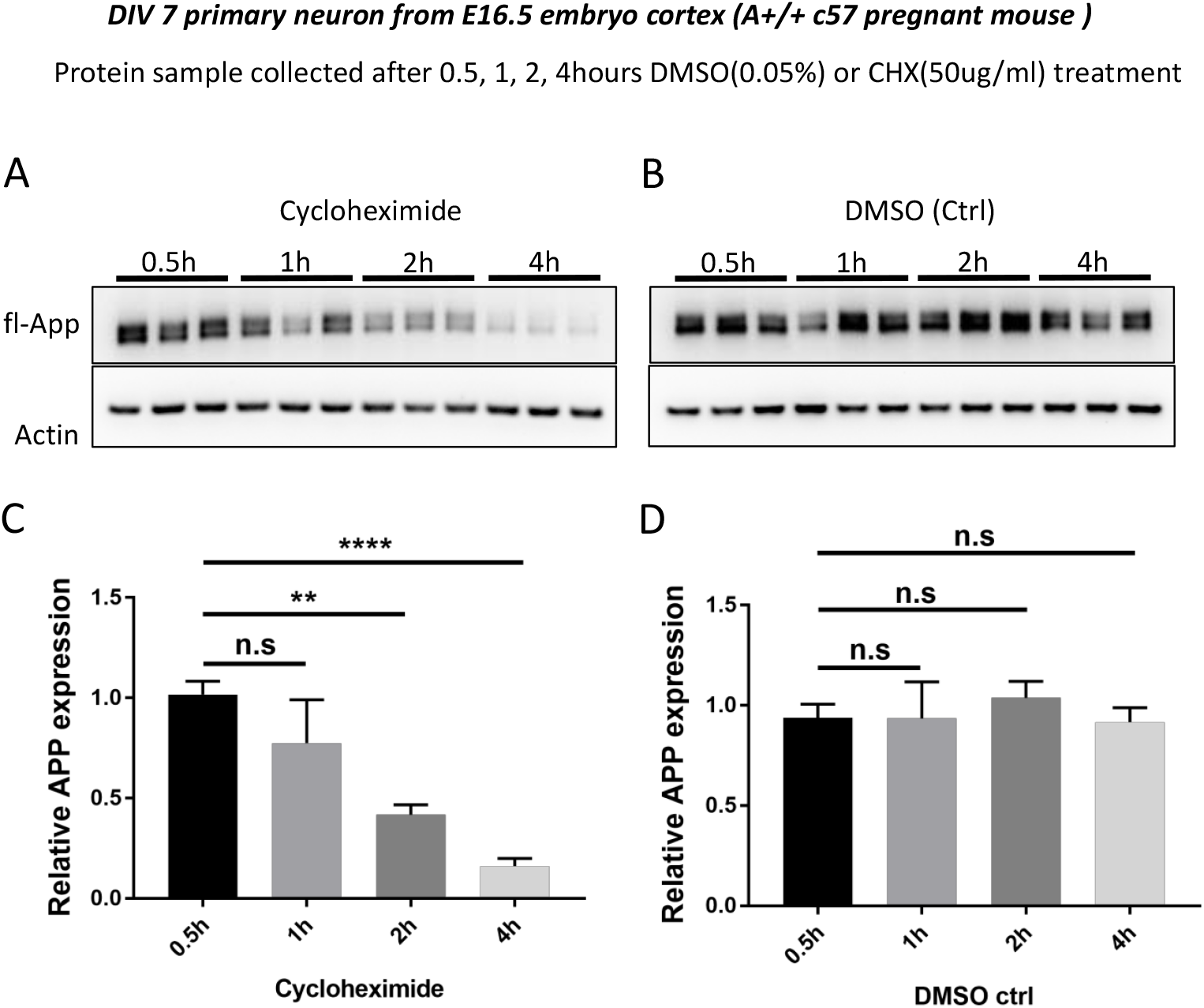
Rapid turn over of fl-mAPP in culture mouse primary cortical neurons. (A-B) Time course (05h, 1h, 2h, 4h) of fl-mAPP expression after Cycloheximide(50ug/ml) or DMSO(0.05%) treatment at DIV7. (C-D) Quantification of fl-mAPP expression after Cycloheximide or DMSO treatment. Bars represent the mean±s.e.m. for at least two independent experiments. **P<0.01, ****P<0.0001

**Figure S4:**
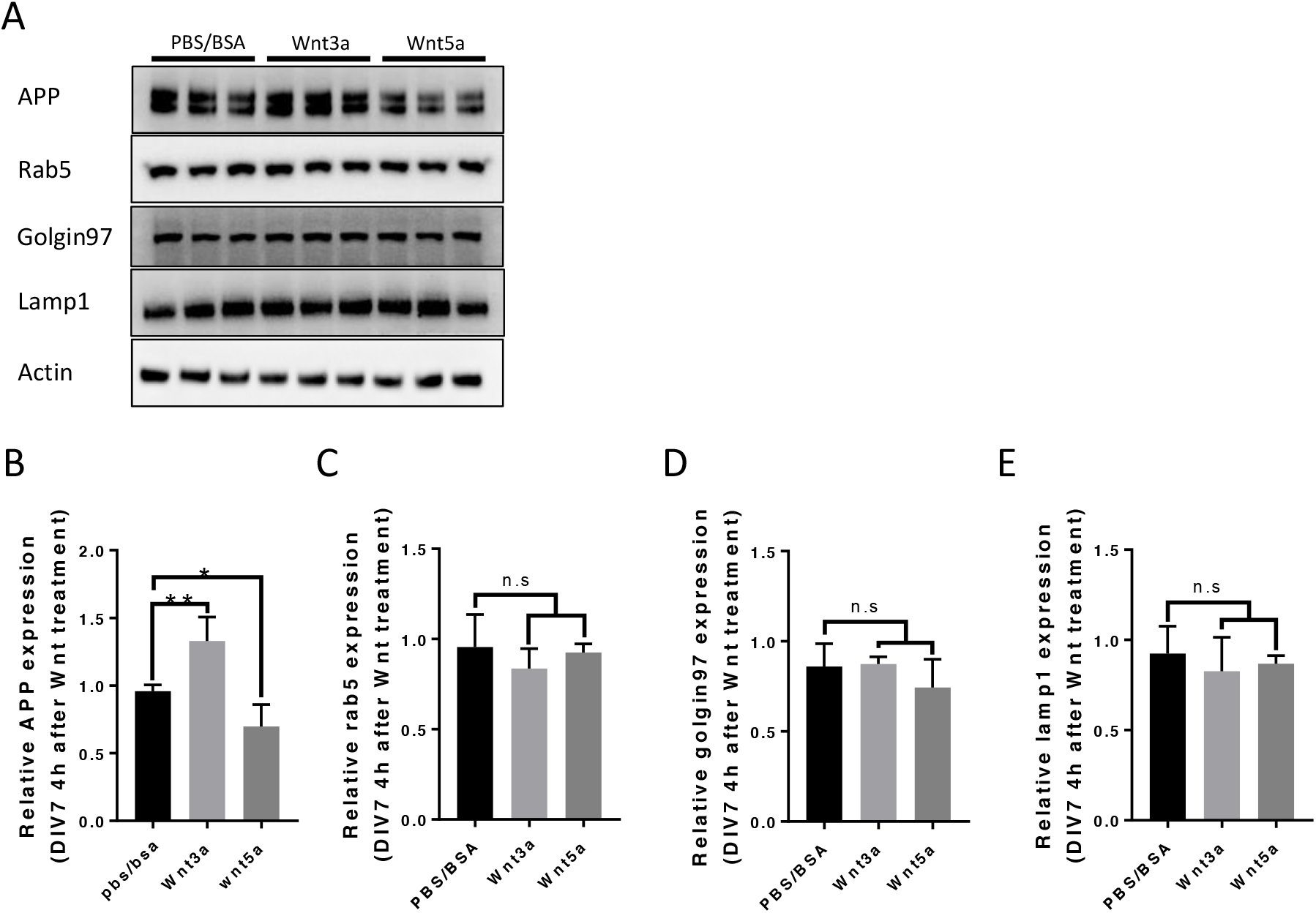
Rab5 Golgin97 and Lamp1 expression after Wnt3a/5a treatment. (A) Protein expression of fl-mAPP Rab5 Golgin97 and Lamp1 after 4hours Wnt3a/5a treatment at DIV7. (B-E) Quantification of fl-mAPP Rab5 Golgin97 and Lamp1 after Wnt3a/5a treatment. Bars represent the mean±s.e.m. for at least two independent experiments. *P<0.05, **P<0.01

**Figure S5:**
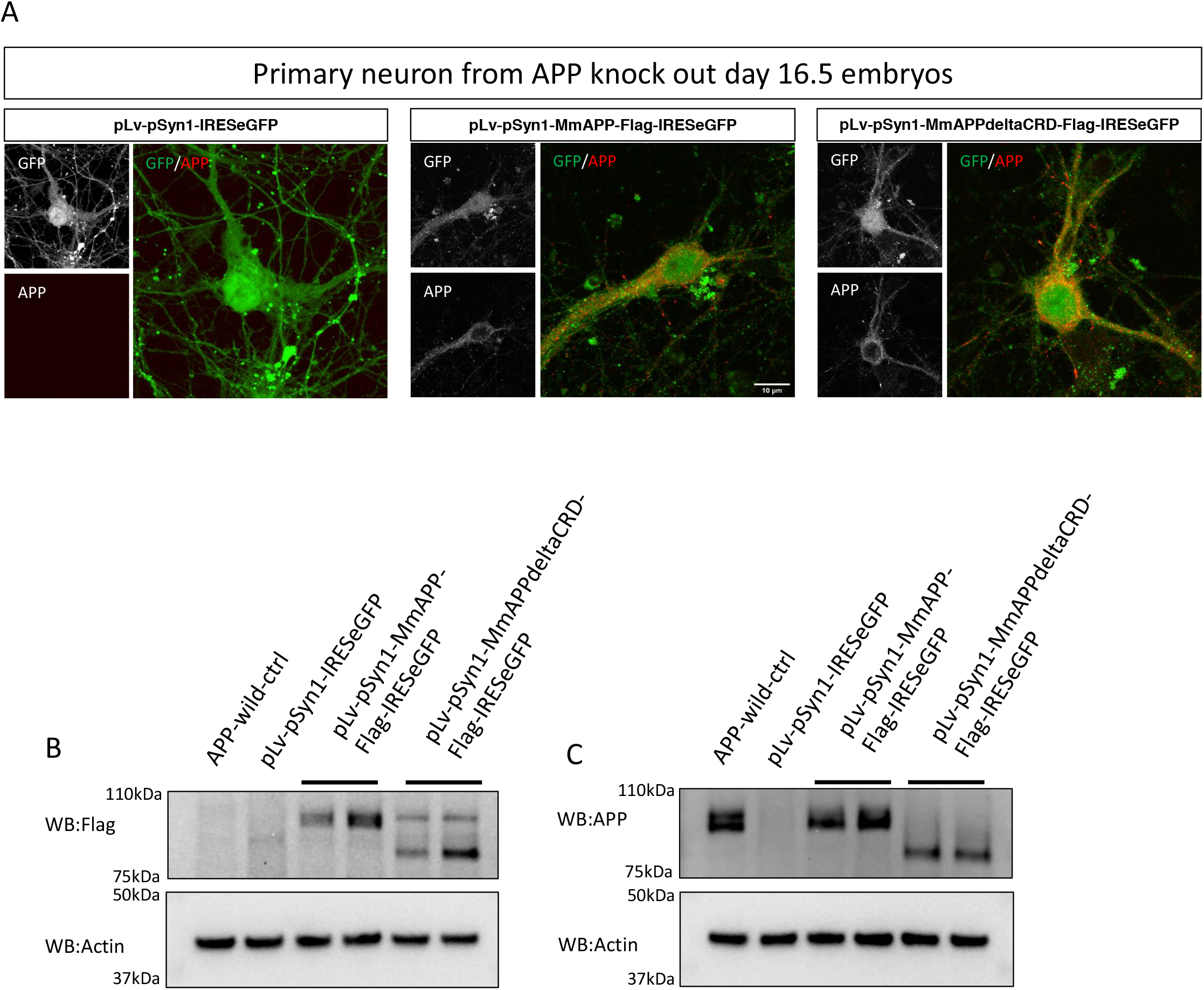
Lenti-virus induced exogenous mAPP expressed in mAPP knock out primary cortical neuron. (A) Figures in middle and left panel shows mAPP protein induced by lenti-virus pLv-pSyn1-mAPP-Flag-IRESeGFP or pLv-pSyn1-mAPP Δ CRD-Flag-IRESeGFP respectively could be detected by immunoflouresouce, right panel is a negative control which transducted with pLv-pSyn1-IRESeGFP. (B-C) Exogenous mAPP could be detected by western blot with anti-flag or anti-APP antibody. scale bar = 10um

**Figure S6:**
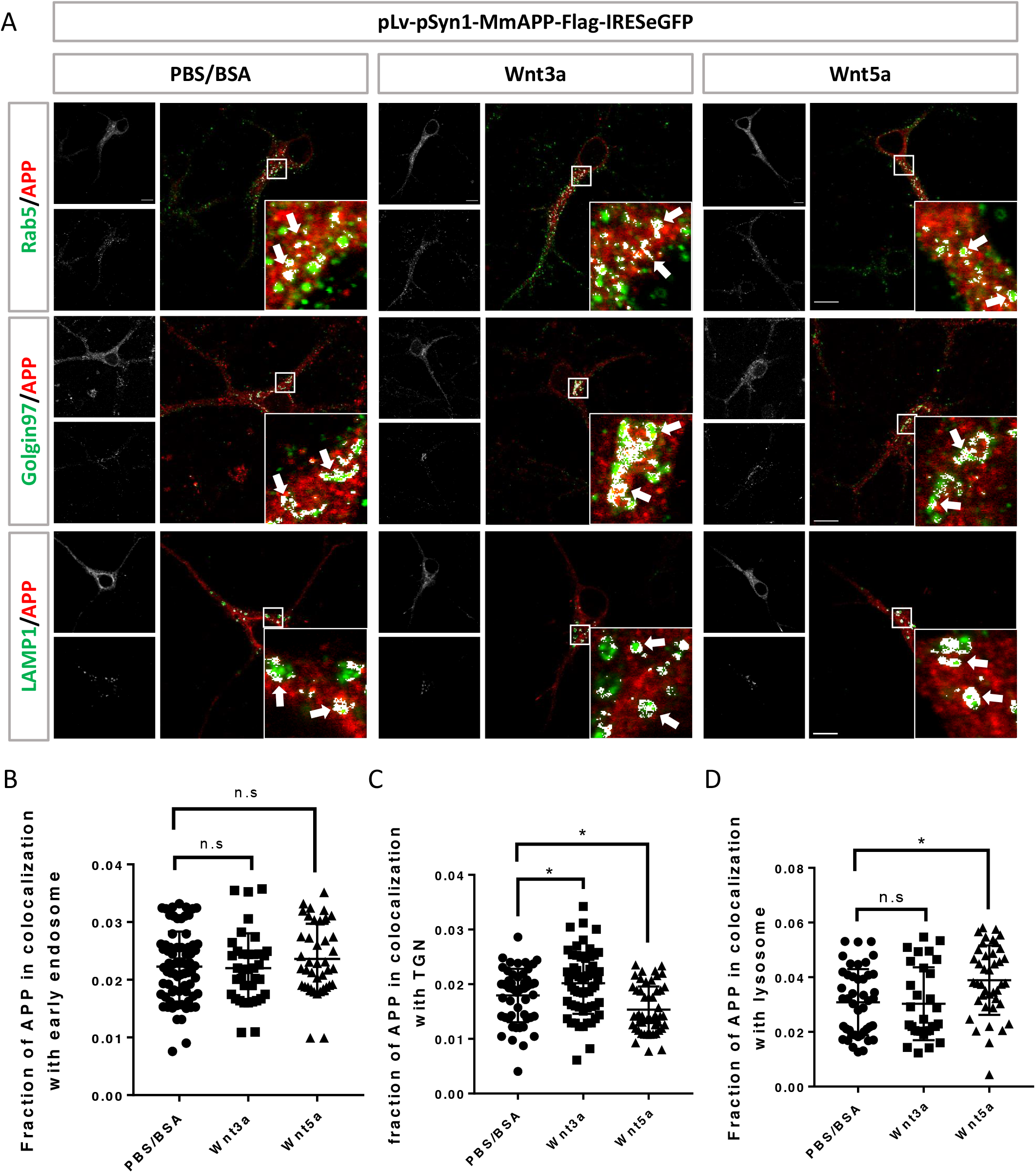
Lenti-virus induced exogenous interact with Wnts in mAPP knock out primary cortical neuron. (A-D) Interaction between exogenous wild type mAPP with Wnts (A) location of exogenous mAPP in APP knock-out primary cortical neurons After 4 hour Wnt3a or Wnt5a treatment. Immunofluorescence using antibody app with rab5(early endosome marker) golgin97(TGN marker) or lamp1(lysosome marker) to reveal mAPP location in different intracellular compartment, the inset fig with White arrow is a high zoom in of the area in white box, arrow indicate the overlap of mAPP with relative cellular compartment. (B-D) quantification of the overlap between mAPP and early endosome TGN or lysosome respectively after Wnt3a or Wnt5a treatment. *P<0.05. scale bar = 10um.

**Figure S7:**
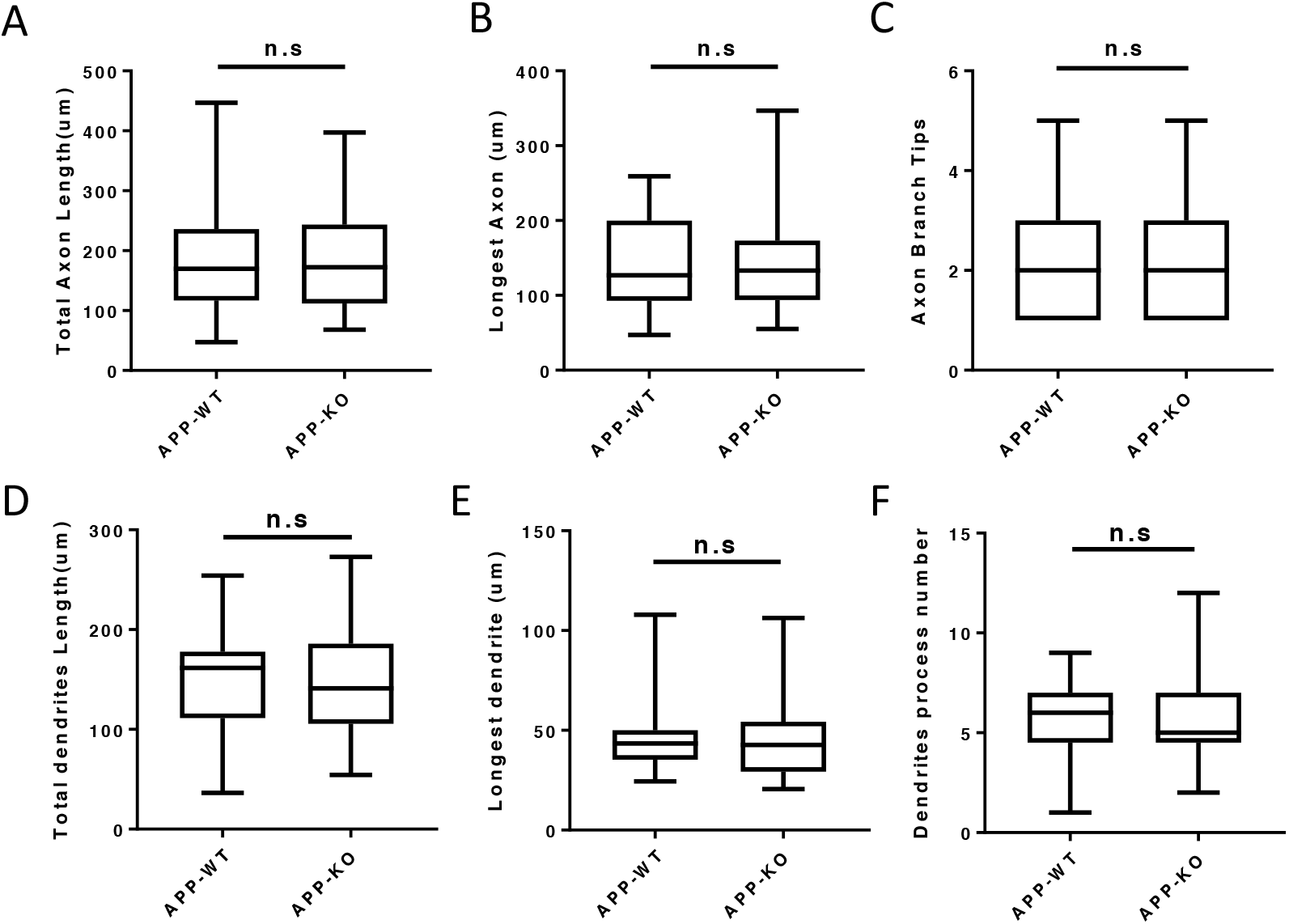
Neurite outgrowth is unaffected in APP knock out neurons at Div2. Loss of mAPP barely affect neurite outgrowth at DIV2. (A-C) Quantification of total axon length, longest axon length and axon branch tips from DIV2 cultured primary cortical neuron. (D-F) Quantification of total dendrite length, longest dendrite length and dendrite branch tips from Div2 cultured primary cortical neuron. Bars represent the mean±s.e.m. for each group, at least two independent experiments, n=50-60.

**Figure S8:**
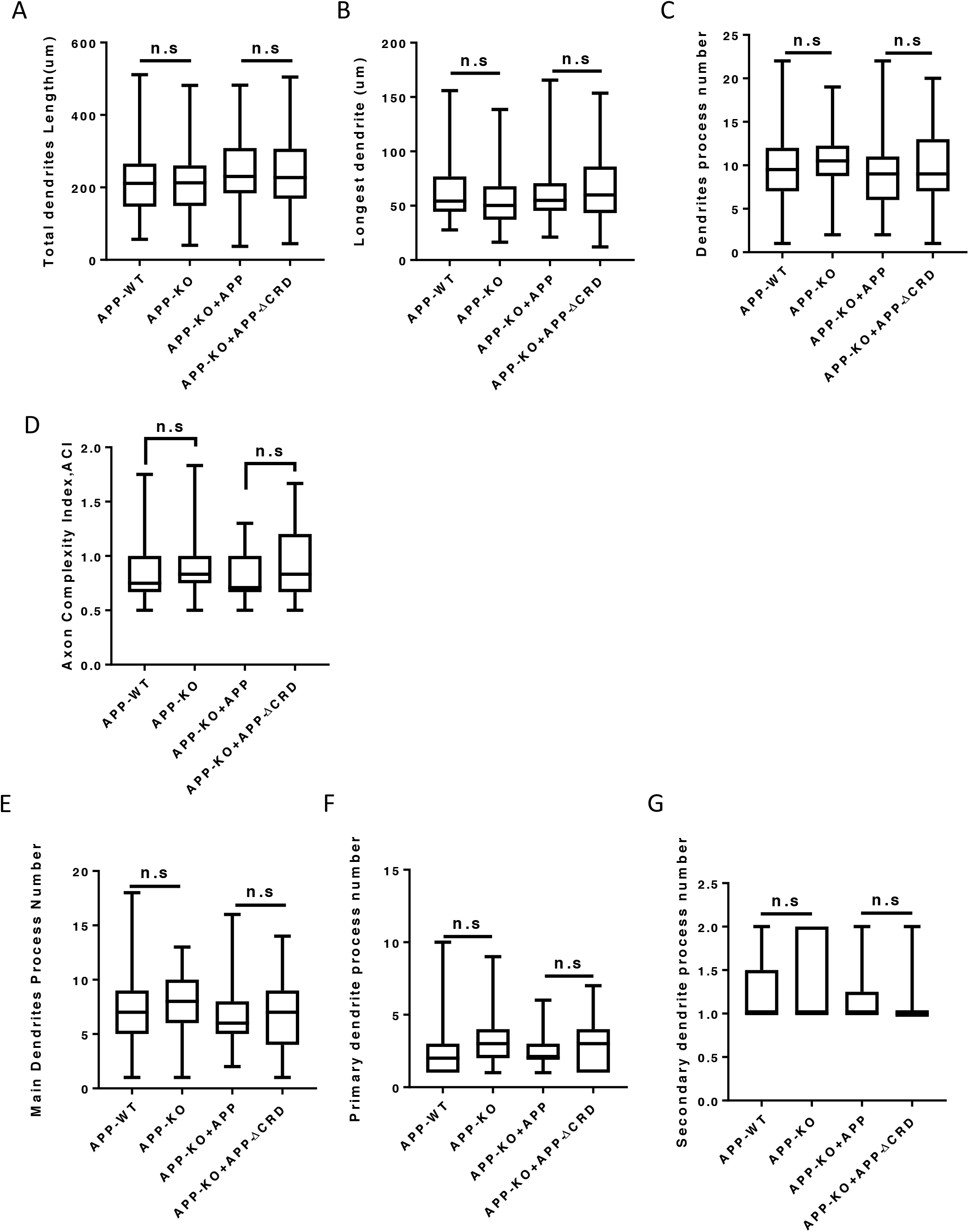
Analysis of dendritic outgrowth and axon complexity index at DIV3. APP mutant barely changes dendrite development at DIV3. (A-C) Quantification of (A) total dendrite length, (B) longest dendrite length and (C) dendrite branch tips from DIV3 cultured primary cortical neuron. (D) Axon complexity Index (ACI) analysis of axon in DIV3 of APP-WT APP-KO and APP-KO with rescue. (E-G) Quantification of (E) main dendrite process, (F) primary and (G) secondary dendrite process numbers.

**Figure S9:**
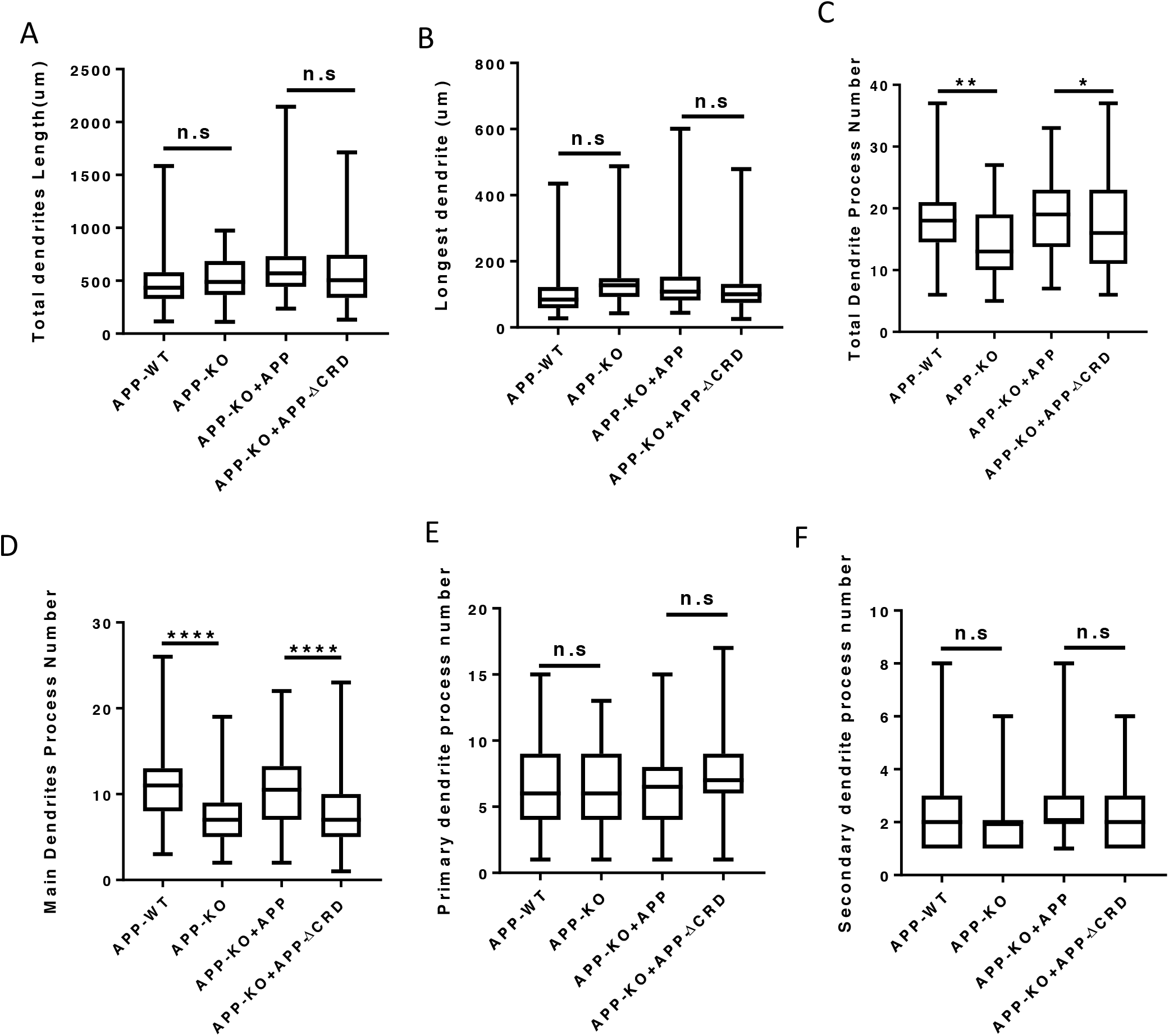
Outgrowth and Complexity analysis of neurite at DIV7. (A-C) Quantification of (A) total dendrite length, (B) longest dendrite length and (C) dendrite branch tips from DIV7 cultured primary cortical neuron. (D-F) Quantification of (D) main dendrite process, (E)primary and (F)secondary dendrite process numbers from DIV7 cultured primary cortical neuron. ****P<0.0001

**Figure S10:**
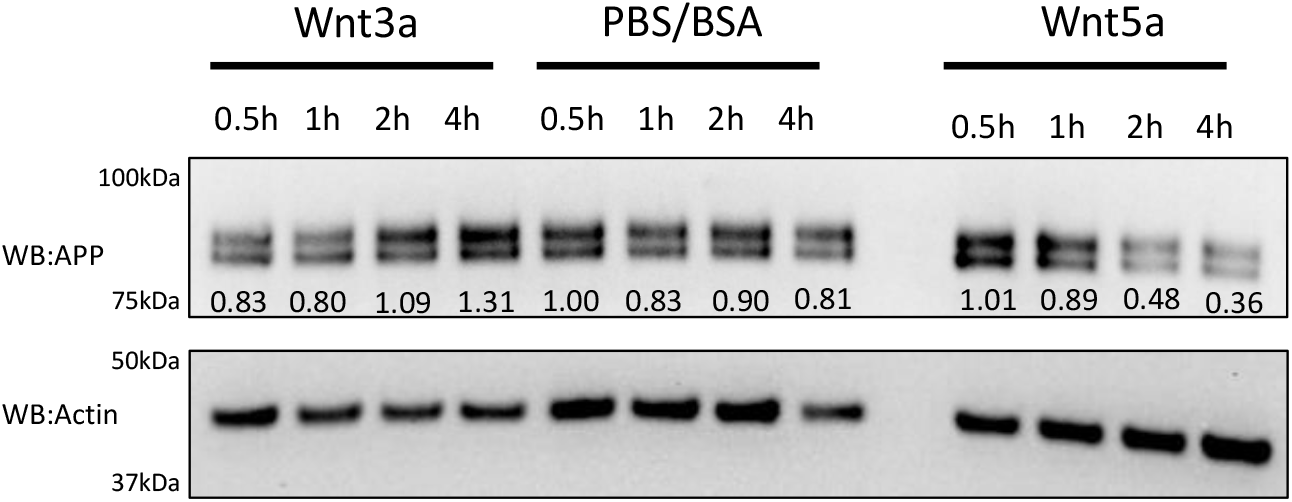
Time course of fl-mAPP after Wnt3a/5a treatment. Representative western blot for the time course (05h, 1h, 2h, 4h) of fl-mAPP expression after PBS/BSA (ctrl) and Wnt3a/5a treatment at DIV7. Relative expression value of the protein bands normalized to the respective actin were also shown

